# Inferring population histories for ancient genomes using genome-wide genealogies

**DOI:** 10.1101/2021.02.17.431573

**Authors:** Leo Speidel, Lara Cassidy, Robert W. Davies, Garrett Hellenthal, Pontus Skoglund, Simon R. Myers

## Abstract

Ancient genomes anchor genealogies in directly observed historical genetic variation, and contextualise ancestral lineages with archaeological insights into their geography and lifestyles. We introduce an extension of the *Relate* algorithm to incorporate ancient genomes and reconstruct the joint genealogies of 14 previously published high-coverage ancients and 278 present-day individuals of the Simons Genome Diversity Project. As the majority of ancient genomes are of lower coverage and cannot be directly built into genealogies, we additionally present a fast and scalable method, *Colate,* for inferring coalescence rates between low-coverage genomes without requiring phasing or imputation. Our method leverages sharing patterns of mutations dated using a genealogy to construct a likelihood, which is maximised using an expectation-maximisation algorithm. We apply *Colate* to 430 ancient human shotgun genomes of >0.5x mean coverage. Using *Relate* and *Colate,* we characterise dynamic population structure, such as repeated partial population replacements in Ireland, and gene-flow between early farmer and European hunter-gatherer groups. We further show that the previously reported increase in the TCC/TTC mutation rate, which is strongest in West Eurasians among present-day people, was already widespread across West Eurasia in the Late Glacial Period ~10k - 15k years ago, is strongest in Neolithic and Anatolian farmers, and is remarkably well predicted by the coalescence rates between other genomes and a 10,000-year-old Anatolian individual. This suggests that the driver of this signal originated in ancestors of ancient Anatolia >14k years ago, but was already absent by the Mesolithic and may indicate a genetic link between the Near East and European hunter-gatherer groups in the Late Paleolithic.

## 1 Introduction

Genetic variation is shaped through evolutionary processes acting on our genomes over hundreds of millennia, including past migrations, isolation by distance, mutation or recombination rate changes, and natural selection. Such events are reflected in the genealogical trees that relate individuals back in time. While these are unobserved, recent advances have made their reconstruction from genetic variation data feasible for many thousands of individuals and have enabled powerful inferences of our genetic past (Rasmussen et al. 2014; Speidel et al. 2019; Kelleher et al. 2019).

Ancient genomes provide a direct snapshot of historical genetic variation, and so add substantial information compared to genealogies built only from modern-day samples. We introduce an extension to the *Relate* algorithm to enable the incorporation of such non-contemporary samples. We use this approach to reconstruct joint genealogies of the Simon’s Genome Diversity Project (SGDP) dataset (Mallick et al. 2016) and 14 previously published high-coverage ancient humans (Cassidy et al. 2020; Broushaki et al. 2016; Jones et al. 2015; Sikora et al. 2017; 2019; Gallego-Llorente et al. 2015; Lazaridis et al. 2014; Fu et al. 2014; Günther et al. 2018; de Barros Damgaard et al. 2018). These genealogies are able to capture the shared population histories of present-day and ancient humans, and could also be applied in other species. In particular, they allow identification of inbreeding, population size estimation and estimation of coalescence rates between individuals, analysis of the age and spread of individual mutations, and in future might be used to infer natural selection (Speidel et al. 2019).

The joint inference of genealogies for ancients and moderns currently requires accurate genotypes, which is not possible for the majority of ancient human genomes which are of lower coverage. One central set of questions for such samples involve estimation of their joint genetic history: their relationships with one another, and other samples, through time, reflected in their varying coalescence rates through time. These coalescence rates can be estimated using a number of methods (H. Li and Durbin 2011; Schiffels and Durbin 2014; Terhorst, Kamm, and Song 2017; Gutenkunst et al. 2009; Kamm et al. 2020), as well as our updated *Relate* approach, but to date none of these have been designed to work for low-coverage genomes. We have therefore developed a fast and scalable method, *Colate,* for inferring coalescence rates between low-coverage genomes without requiring phasing or imputation. *Colate* leverages age distributions of mutations from a *Relate*-inferred genealogy to construct a likelihood that summarises sharing patterns of mutations through time, which we maximise using an Expectation-Maximisation (EM) algorithm. The method can calculate coalescence rates between any number of samples and scales linearly in sample size and genome lengths; *Colate* requires only a constant runtime of typically 5 seconds for the EM step after parsing the data (**Methods**).

We apply *Colate* to 430 genomes of >0.5x coverage spanning the late Paleolithic, Mesolithic, Neolithic, and more recent epochs across many regions outside Africa (SI Table). Using *Colate*-inferred coalescence rates, as well as our *Relate* results for higher-coverage genomes, we trace genetic structure evolving through time. Among other findings, we readily identify genetic clusters corresponding to HGs, Neolithic farmers, and the Bronze age in Europe, and map out the coalescence rates of modern humans worldwide with these ancient samples. We show that these indicate localised structure, and characterise dramatic population replacements in Ireland within the space of 3,000 years, as well as varying gene flow between HGs and Neolithic farmers across Europe, which is more widespread than previously identified.

Finally, we leverage our *Relate*-inferred genealogies and *Colate*-inferred coalescence rates to quantify the previously reported but unexplained elevation in TCC to TTC mutation rate (K Harris 2015) in all SGDP individuals and 161 ancient individuals of >2x mean coverage, providing a finer-scale geographic and temporal mapping of this signal than previously available. We show that the signal has a remarkable 96% correlation with coalescence rates to an early Anatolian farmers from the pre-pottery Neolithic (Kılınç et al. 2016), is absent in samples from >34,000YBP but was already widespread among HGs in Late Glacial West Eurasia, and shows no increase in strength over the last 10,000 years, suggesting that the driver for this excess was extinct by the Late Mesolithic. This strong localisation of the signal in both time and space suggests either a genetic cause, or a somehow tightly focussed environmental cause. Moreover, we hypothesise that these excess TCC/TTC mutations spread via gene flow through ancestors of ancient Anatolia into HG groups across Western Eurasia before the expansion of farming, perhaps associated with a link between the Near East and Late Upper Paleolithic Europe that started with the Bølling-Allerød interstadial warming period (Fu et al. 2016).

## 2 Results

### 2.1 Extending *Relate* to work with non-contemporary samples

We extend our previously developed method, *Relate*, for inference of genealogical trees genome-wide for large sample sizes (Speidel et al. 2019) to work with ancient genomes (Supplementary Information). A key aspect of non-contemporary samples is that these impose hard constraints on the ages of coalescence events. Our updated tree builder restricts which lineages can coalesce by assigning a preliminary date to each coalescence event and only allows coalescences of non-contemporary samples with lineages that predate its age. Branch lengths are sampled using a Markov-Chain Monte Carlo sampler, with modified proposal distributions to allow for non-contemporary samples. As before, we sample branch lengths from a posterior distribution that fixes tree topology and combines the likelihood of observing a certain number of mutations on a branch and a coalescent prior with piecewise-constant effective population sizes through time.

### 2.2 Inferring coalescence rates for low-coverage genomes using *Colate*

*Colate* calculates coalescence rates between a set of “target” and a set of “reference” chromosomes by leveraging mutations dated using an inferred genealogy; this genealogy may (or may not) have overlapping samples with the target and reference chromosome sets (Figure 1, **Methods** and Supplementary Information). Both the target and reference chromosomes may be specified as VCF files containing genotypes, or as BAM files containing reference-aligned reads. The latter is particularly useful for low-coverage sequencing data, where accurate genotype calling is not possible. For ancient genomes, we specify a sampling date. In practise, we often specify two different individuals as the target and reference, and obtain the coalescence rates between this pair, though it would be possible to pool information.

**Figure 1.**
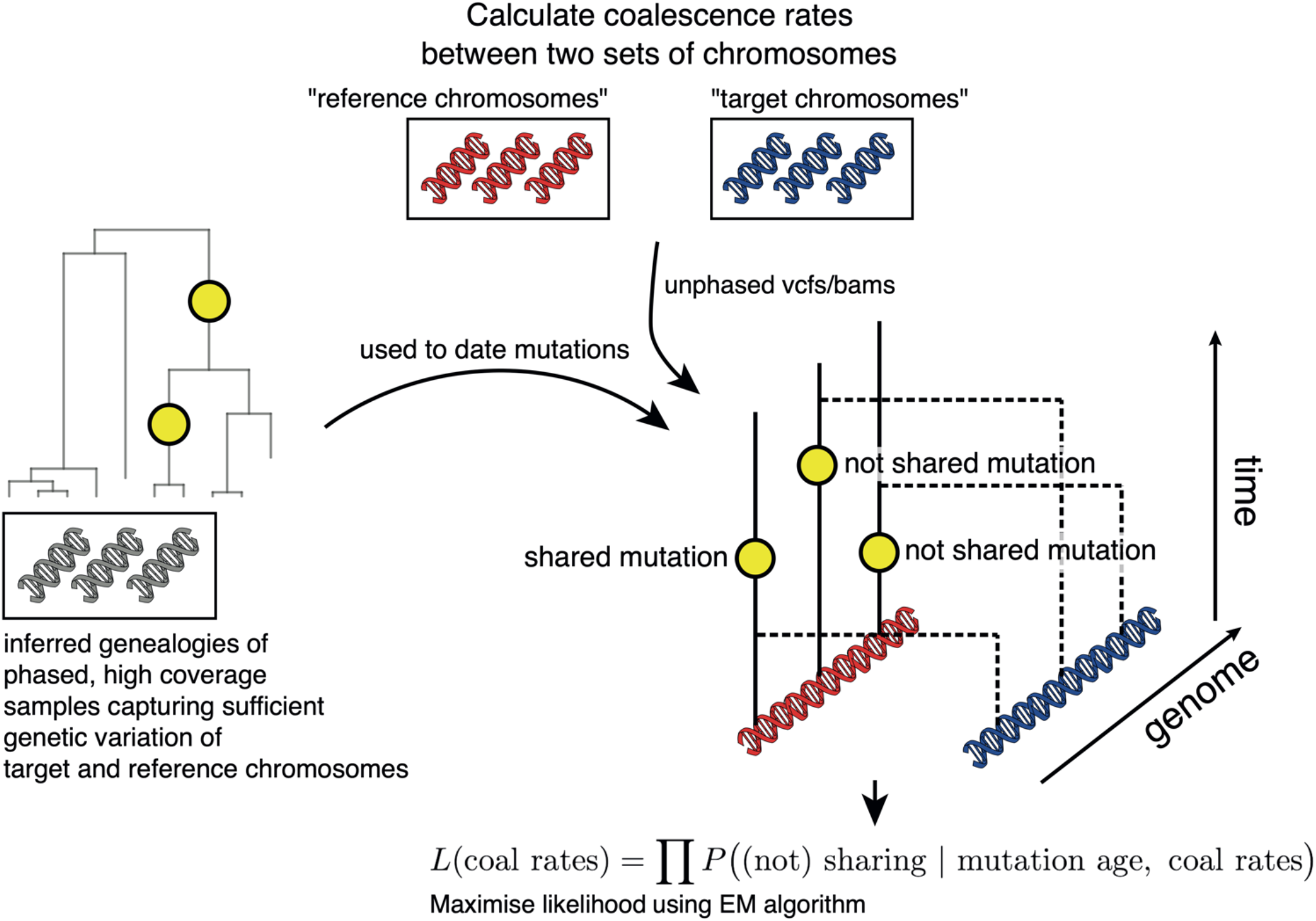
*Colate* calculates coalescence rates between two sets of chromosomes, labelled target and reference (main text). The method proceeds by recording for each mutation carried by a reference chromosome, whether it is shared in the target chromosomes. This information is summarised in a likelihood, which is constructed by multiplying over SNPs, such that no phase information is required. Whenever more than one chromosome is available at any given site, we multiply across chromosomes. The likelihood is maximised using an expectation-maximisation algorithm.

The *Colate* likelihood uses as input data whether each mutation carried by a reference chromosome is shared, or not shared, with a target chromosome. Sharing indicates that coalescence between the two chromosomes happened more recently than the age of this mutation, whereas non-sharing indicates that coalescence happened further in the past, assuming each mutation occurs only once (the infinite-sites model), and so an exact likelihood can be calculated, given coalescence rates between the sample sets (**Methods**). We multiply this likelihood across sites and therefore do not require genomes to be phased; in low-coverage data, we additionally multiply across pairs of reads. This likelihood is then maximised using an expectation-maximisation (EM) algorithm (**Methods,** Supplementary Information). Our implementation reduces computation time by using a discrete time grid to record sharing and non-sharing of mutations through time, reducing the computation time of the EM algorithm. As a result, computation time is independent of sample size and genome lengths once the data is parsed, and typically takes around 5 seconds (~40 seconds including parsing the data, Supplementary Figure 1).

We demonstrate high accuracy of *Colate* and *Relate*-inferred coalescence rates using the stdpopsim package (Adrion et al. 2020), on simulated data following a zigzag demographic history (Supplementary Figure 2) as well as a multi-population model of ancient Eurasia, which was fitted using real human genomes (Kamm et al. 2020) (Figure 2**a**) (**Methods;** also see (Speidel et al. 2019) for comparison of *Relate* to other methods). We further evaluate *Colate*’s performance on low-coverage sequencing data by downsampling high-coverage genomes of the 1000 Genomes Project (The 1000 Genomes Project Consortium 2015), and find that although uncertainty increases as coverage decreases, *Colate* recovers meaningful coalescence rate estimates even between a sequence of 0.01x mean coverage and high-coverage sequences specified as a VCF (Figure 2**b**), or between two low coverage sequences of 0.1x mean coverage (Supplementary Figure 3).

**Figure 2.**
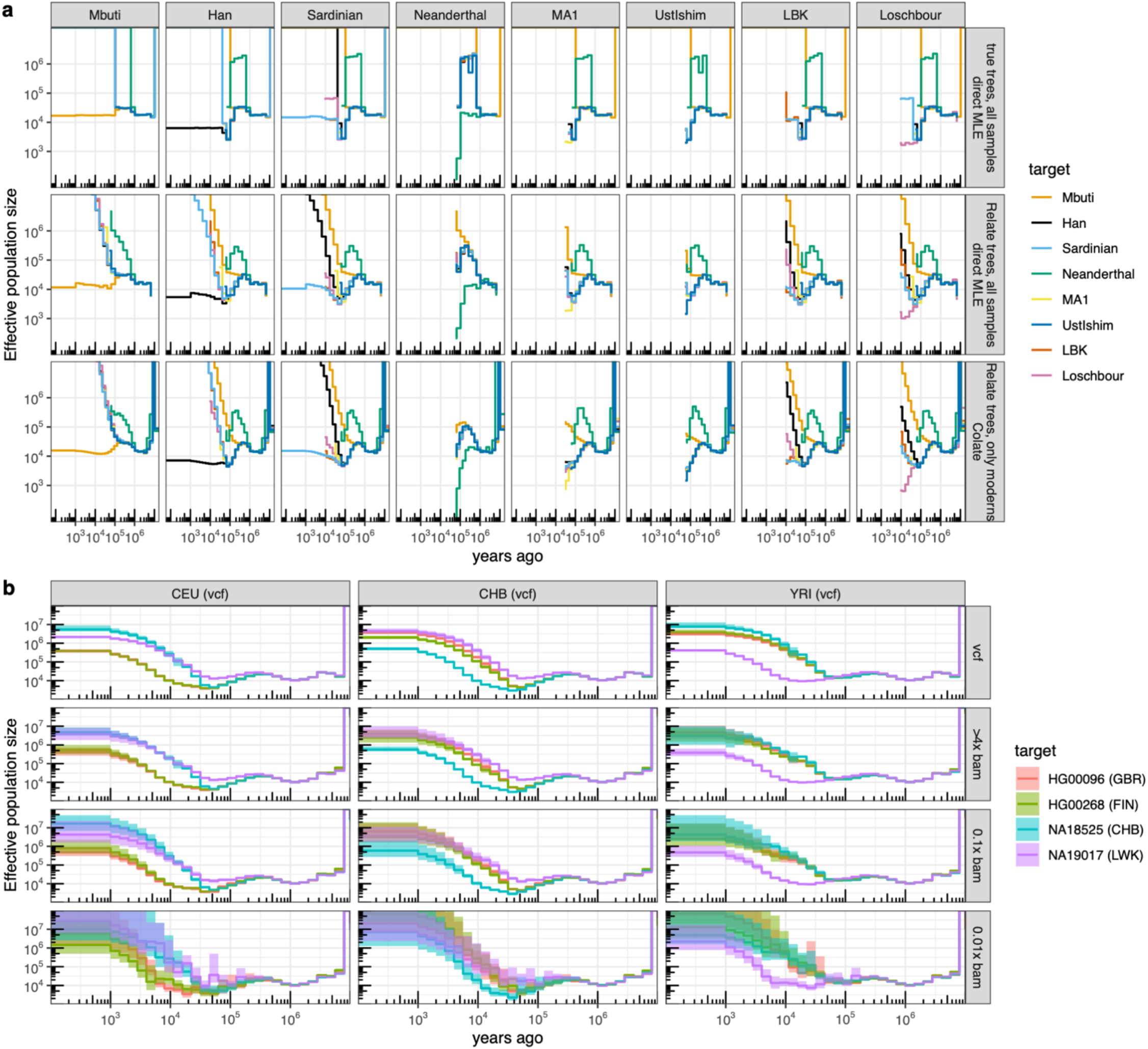
**a**, Simulation emulating real human groups, including three modern human groups (Mbuti, Han, and Sardinian) with 100 haploid sequences each, and five diploid ancient genomes. We calculated coalescence rates between groups using true genealogical trees of all samples (true trees; direct MLE), *Relate* trees of all samples (*Relate* trees; direct MLE), as well as *Colate*, where the reference genealogy included all modern human groups but not the ancients. For the direct MLEs, coalescence rates are symmetric with respect to target and reference group assignment; for *Colate*, each panel corresponds to a fixed reference group, with different coloured lines showing different target groups. **b**, *Colate*-inferred coalescence rates between four 1000 Genomes Project samples (HG0096, HG00268, NA18525, NA19017) and the remaining 1000 Genomes samples in groups CEU, CHB, and YRI. We calculate coalescence rates where the target samples are given as genotype data (VCF), as well as reference-aligned read data downsampled to 4x, 0.1x, and 0.01x mean coverage. Confidence intervals are constructed using 100 block bootstrap iterations with a block size of 20Mb.

### 2.3 *Relate* and *Colate* applied to 278 SGDP moderns and 430 ancients

We inferred joint genealogies of 278 modern-day individuals of the Simons Genome Diversity Project and 14 previously published high coverage ancients of >8x mean coverage, which we collectively rephase using Shapeit4 (Delaneau et al. 2019) and the 1000 Genomes Project reference panel (**Methods**). Tree topologies were constructed using all mutations except CpG dinucleotides, but branch length inference used transversions only, to avoid confounding due to deamination errors in the ancient genome sequences (**Methods**). Additionally, we estimate pairwise-coalescence rates for 430 ancient individuals of >0.5x mean sequencing coverage using *Colate* (SI Table). For *Colate*, we use a *Relate*-inferred genealogy for the SGDP samples to date mutations, where we sampled one haplotype from each individual to remove the effects of recent inbreeding and restrict to transversions (**Methods**).

### 2.4 PCA on *Colate*-inferred coalescence rates captures dynamic population structure

*Colate*-inferred coalescence rates demonstrate intricate relationships that vary geographically and through time and manifest vast migrations and, in places, repeated population replacements (Figure 3**a**,**b**). In the recent past (0-15KY), populations are separated based on both geography and sample age (Figure 3**a**,**b**): there are extremely low coalescence rates between continental regions (excepting W. Eurasia, Central Asia, and Siberia, which show patterns indicating migration). Taking samples from Ireland as one example (Figure 3**b**), previous work has indicated repeated partial or complete population replacements, first of ancestral hunter-gatherers by Neolithic farmers, and then in the Bronze age by migrants related to people from the Eastern steppe (Cassidy et al. 2016). Using *Colate,* the earliest Irish samples have highest coalescence rates with, and similar relatedness to other groups as, West European hunter-gatherers (e.g. Loschbour). Neolithic Irish samples show much lower affinity to these hunter-gatherers, but are closely similar to other European farmers (e.g. LBK, an early farmer from Germany). Bronze age Irish samples again show more similarity to hunter gatherers, but now *Eastern* European huntergatherers (and other Eastern European groups), and in this and other respects they resemble the Yamnaya, a possible source group (Figure 3**b**); however they retain some farmer-like haplotypes not present in the Yamnaya sample. Comparing across the whole dataset, we observe that Irish ancient genomes are closest to other Irish ancients from within the same time period (Supplementary Figure 4, 5). This implies that finer scale, regional stratification existed within the HGs, Neolithic farmers, and Bronze age samples, but there is no clear evidence of continuity across periods, suggesting this arose independently repeatedly. We also identify clear substructure among European HGs, consistent with previous findings (Lazaridis et al. 2014) and pairwise F2 statistics (Supplementary Figure 6); this structure corresponds to a divide of Western, Eastern, Scandinavian, and Caucasus HGs among our samples in Europe.

**Figure 3.**
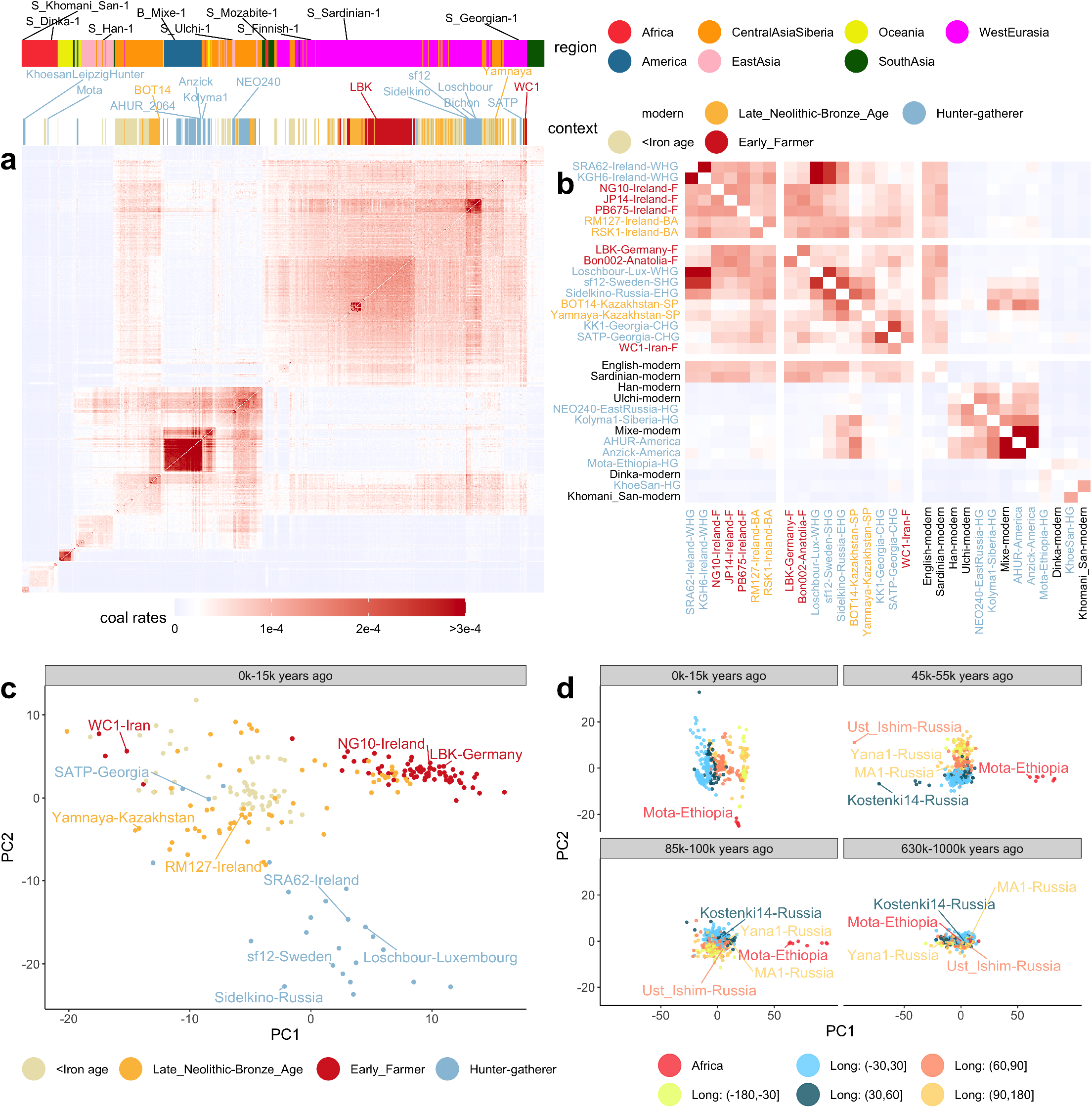
**a**, Matrix of pairwise coalescence rates of all SGDP individuals and ancients in epoch 0 to 15,000 years before present (YBP), calculated using *Colate.* **b**, Subset of samples shown in **a.** Sample names are coloured by context. Abbreviations in sample names are WHG: Western hunter-gatherer, SHG: Scandinavian hunter-gatherer, EHG: Eastern hunter-gatherer, CHG: Caucasus hunter-gatherer, F: farmer, BA: Bronze Age, SP: Steppe Pastoralists **c**, Principal component analysis (PCA) on pairwise coalescence rates of ancient individuals in epoch 0 - 15,000 YBP, coloured by context. **d**, PCA on pairwise coalescence rates for four epochs, coloured by Longitude outside Africa. In all PCAs, we standardised columns in each matrix of coalescence rates and applied the R function prcomp to calculate PCs.

One approach to visualise the diverse signals in these data is to adapt the widely used PCA approach, but now using coalescence rates within particular epochs (Figure 3**c**,**d** show the first two PCs for selected epochs). Structure is not seen in the deep past (>630k years before present (YBP)) but in distinct epochs we observe separation first of African (e.g., Mota) and non-African individuals, and by 45-55k YBP, a separation between West and East Eurasians, as well as a stronger split with Ust’-Ishim (Fu et al. 2014), a 45k-year-old Siberian individual who also appears slightly closer to East Eurasians compared to later European samples, such as Kostenki14 (Seguin-Orlando et al. 2014) and Sunghir3 (Sikora et al. 2017), who are closer to West Eurasians. In the most recent epoch (0-15k YBP), our PCA mirrors geography globally (Novembre et al. 2008), but reflects different ancestries more regionally; for instance, we detect three clusters, corresponding to Mesolithic HGs, Neolithic farmers, and Bronze/Iron age individuals in Europe (Figure 3**c**). The Bronze age cluster falls closer to Steppe Pastoralists from the Pontic-Caspian Steppe (e.g., Yamnaya), consistent with previously reported gene flow from this region into Bronze age Europe (Haak et al. 2015; Allentoft et al. 2015). Overall, these inferences seem in strong agreement, across time and space, with previous specific analyses of these samples.

### 2.5 Relationship of European hunter-gatherer groups to Neolithic farmers

We assess the ancestry contributions of several potential approximate ancestral sources: early European farmers, Western, Scandinavian, and Caucasus HGs to present-day West Eurasians (Figure 4), by measuring the coalescence rates - quantifying shared ancestry - of modern individuals from each of these groups. As expected, HG ancestry is more localised in present-day Europeans compared to shared ancestry with Neolithic farmers, who arrived to Europe from Anatolia (Haak et al. 2010). We also detect a previously observed South-North cline, with the highest farmer-like ancestry observed in Sardinians (Figure 3**b**), while Western and Scandinavian HG ancestry is highest in northern European groups and Caucasus HG ancestry is concentrated around present-day Georgia (Lazaridis et al. 2014; Skoglund et al. 2012; 2014; Jones et al. 2015).

**Figure 4.**
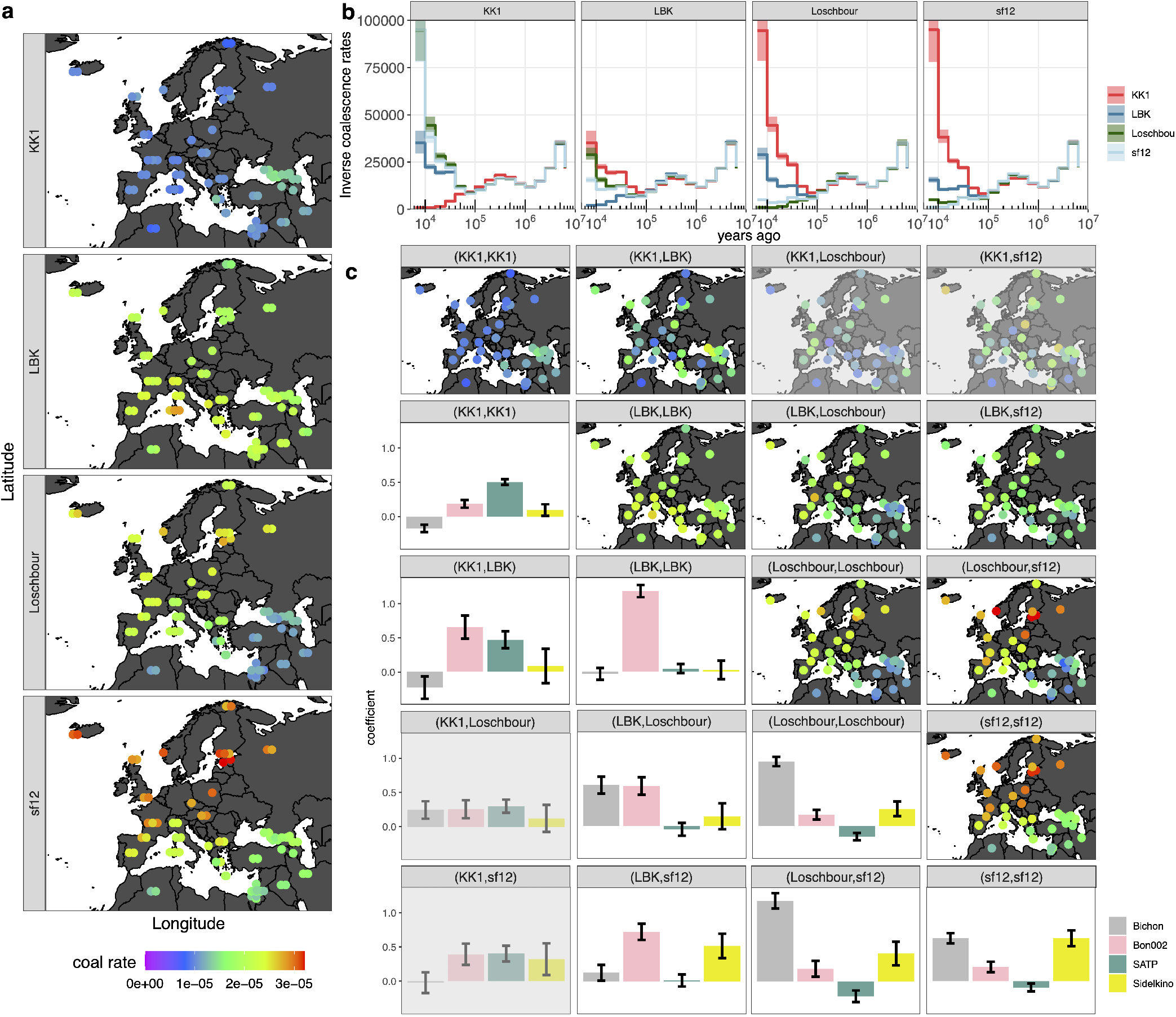
**a**, Map showing *Relate*-inferred coalescence rates of a 9700-year-old Caucasus HG (KK1), 7200-year-old early European farmer (LBK), a nearly 8000-year-old Western hunter-gatherer (Loschbour), and a 9000-year-old Scandinavian HG to SGDP moderns. The coalescence rates shown in the map correspond to the epoch 16k-25k YBP. b, *Relate*-inferred inverse coalescence rates (effective population sizes) for KK1, LBK, Loschbour, and sf12 to themselves and each of the other four individuals. **c**, Maps in top diagonal show *Relate*-inferred coalescence rates of lineages with descendants shown by facet titles to SGDP moderns in same epoch as in **a.** Bottom diagonal shows regression coefficients obtained by regressing coalescence rates (integrated over interval 0-50k YBP) of lineages with descendants given by facet titles to SGDP moderns against *Colate*-inferred coalescence rates (integrated over interval 0-50k YBP) of Bichon (Western HG), Bon002 (Anatolian), SATP (Caucasus HG), Sidelkino (Eastern HG) to SGDP moderns. Panels involving KK1 and Loschbour or sf12 are greyed out, as there is little gene-flow between these groups.

While there is strong evidence for Anatolian farmers partially replacing HG ancestry across Europe in the Neolithic, the deeper relationship of ancestors of these Anatolian farmers to European HGs in the Late Upper Paleolithic is not fully understood. Caucasus HGs have been modelled as forming a clade with European early farmers that is deeply diverged from Western HGs (>27k YBP), with subsequent directional gene flow from Western HGs into Anatolia (Jones et al. 2015). More recent studies have demonstrated that the major ancestral component of Western HGs only became widespread in Europe after 14k YBP and harbours an increased affinity to Anatolian and Caucasus populations, relative to earlier European HGs (Fu et al. 2016), suggesting an expansion from Southeast Europe or the Near East following the Last Glacial Maximum (LGM). To address such questions, we first estimate and characterize overall pairwise coalescence rates among samples. To focus on migration between two groups A and B, we examine lineages that possess descendants in each group as a result of recent shared ancestry, and might therefore represent migrants from one population to another. If recent migration is purely directional from group A into group B, such lineages will always come from group A in the past, and thus have the same coalescence rates as this group (rather than group B).

Initially, using pairwise coalescence rates, we find that Western and Scandinavian HGs form a clade relative to Caucasus HGs (KK1), with almost no recent coalescences observed between these groups. However patterns observed for early farmers (LBK) imply a non-tree-like group relationship involving migration (Figure 4**b**): Caucasus HGs show greater affinity to Neolithic farmers than to Western or Scandinavian HGs in recent epochs, but this is not reciprocated by early farmers who have higher coalescence rates to Western and Scandinavian HGs than to Caucasus HGs.

We therefore characterise lineages ancestral to two haplotypes that coalesced recently (<50k YBP), in the expectation that directional migration would imply that lineages, once they coalesce with the migrating group, will appear similar to lineages ancestral to the migrating group (Figure 4**c**). To gain power, we calculate the coalescence rates of these lineages to each non-African SGDP modern sample, and perform a linear regression against *Colate*-inferred coalescence rates of four individuals representing independent samples from similar, but older groups: ancient Anatolia (Bon002) (Kılınç et al. 2016), Western HGs (Bichon) (Jones et al. 2015), Eastern HGs (Sidelkino) (de Barros Damgaard et al. 2018), and Caucasus HGs (SATP) (Jones et al. 2015) to the same SGDP moderns, to fit these lineages as a mixture of these four potential surrogate source populations. We rescaled *Colate* coalescence rates according to Supplementary Figure 8 to match overall levels of coalescence rates between *Colate* and *Relate.*

Encouragingly, we find that lineages ancestral to the two haplotypes of the same individual (not indicating migration) are well captured by one respective ancestry in our regression in three cases and suggesting these are reasonable surrogates. The exception is the Scandinavian HG (sf12) who we fit as an approximately equal mixture of Eastern and Western HGs, as previously reported (Günther et al. 2018). The highest recent coalescence rates we see are between the Western and Scandinavian HG: recently coalesced lineages between these samples appear very similar to Western HGs (Figure 4**c**), indicating strong directionality of gene-flow, from Western HG into Scandinavia. In contrast, lineages that are ancestral to LBK and any of the other three HGs are fit as a mixture of early Anatolian farmers and the respective HG groups, suggesting gene-flow both into and from ancestors of LBK, though biased in some cases.

### 2.6 Effective population sizes increased from Mesolithic Europe to the present

Effective population sizes calculated within an individual quantify diversity and relatedness of parental genomes. By focussing on the very recent past (<1000 years), we observe a broad spectrum of recent within-individual effective population sizes in SGDP individuals ranging from a few thousand to hundreds of thousands not limited to particular geographical groups (Figure 5**a**, Supplementary Figures 7) and correlating well between *Relate* and *Colate* (Supplementary Figures 8). Haplotypes of individuals with small recent effective population sizes coalesce with each other before coalescing with any other sample for larger proportions of the genome (Figure 5**b**), indicative of longer runs of homozygosity (ROH) in these individuals (Supplementary Figure 9). While global patterns are comparable to previously reported heterozygosity estimates (Mallick et al. 2016), the differences among particular individuals are more pronounced in our analysis, which focusses on very recent time.

**Figure 5.**
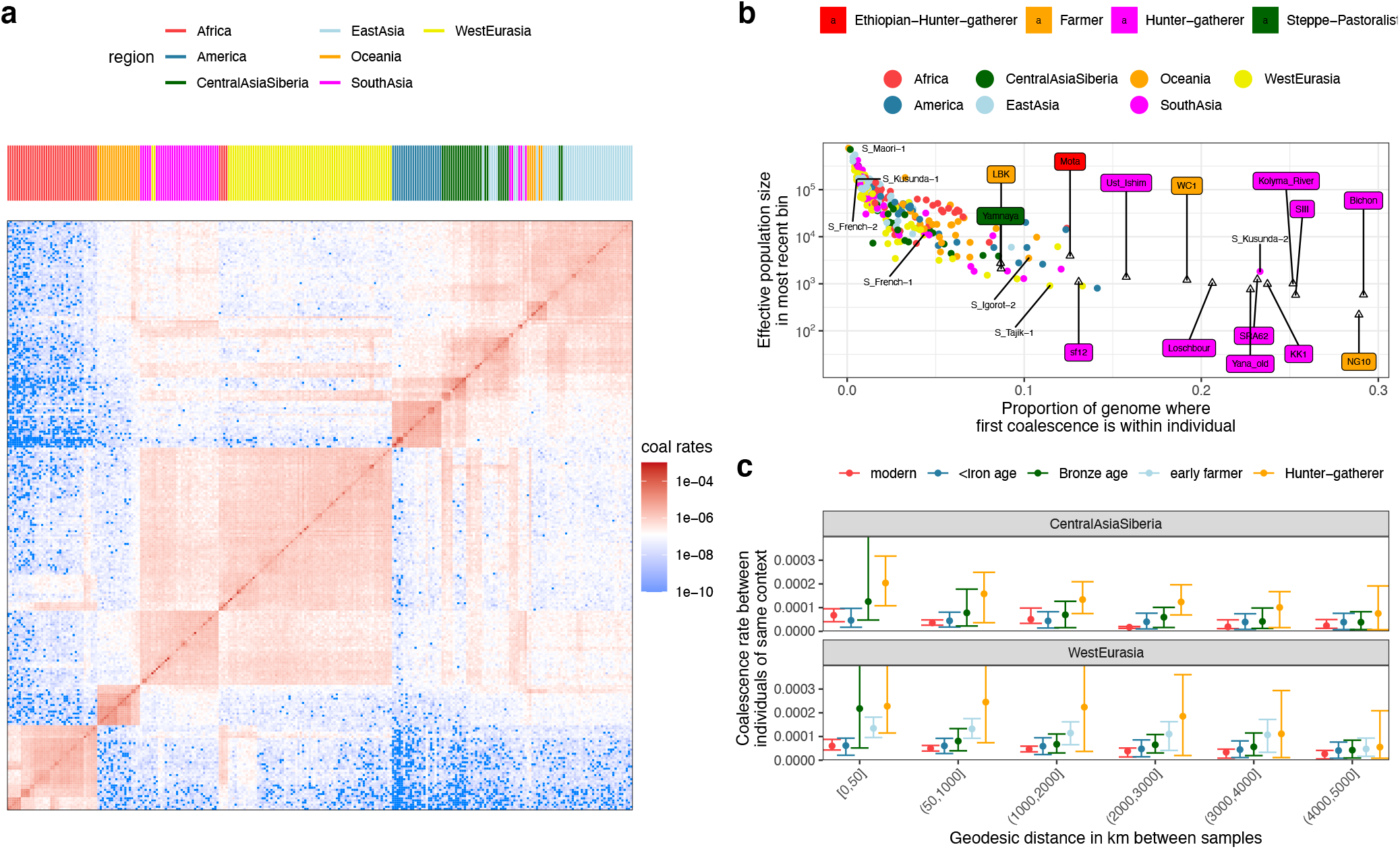
**a**, *Relate*-inferred coalescence rates between SGDP individuals in most recent epoch (0 - 1,000 years BP). **b**, Within individual effective population sizes in the most recent epoch plotted against the proportion of the genome where the first coalescence occurs within the individual. All coalescence rates were calculated using *Relate* trees. **c**, *Colate*-inferred coalescence rates in the most recent epoch (<15k YBP) averaged over pairs of samples grouped by geographic distance and time period. Error bars show the 2.5% and 97.5% percentiles, respectively.

Small recent effective population sizes are also observed in the high coverage ancient genomes and are most pronounced in European Mesolithic HGs, who also tend to coalesce with themselves for larger proportions of the genome, however this may at least in part be driven by increased divergence from other samples, in addition to ROH (Figure 5**b**). The smallest recent effective population size is observed for the NG10 individual, a 5,200-year-old Neolithic individual buried in a Megalithic tomb in Ireland, who was previously identified to be the son of a first-degree incestuous union (Cassidy et al. 2020). We next compared coalescence rates across individuals at increasing geographic distances within Europe, and within Central Asia, in each time period, including only moderns within 500km of an ancient sample (Figure 5**c**). At short distances we observe a clear trend for smaller coalescence rates (larger effective population sizes) towards the present, suggesting strongly increasing local population sizes. At larger distances the relationship is non-monotonic, with coalescence rates not decreasing consistently, implying a trend of increasing migration, countering the larger population sizes. Finally, we see a trend of decreasing similarity with distance, implying local population structure at all times, with the interesting exception of samples more recent than the beginning of the Iron age (yet not modern) in Europe. More widespread sampling is needed to understand this pattern, although this period does overlap e.g., increased mobility during the Roman Empire and the following “migration age” in Europe characterized by widespread movements of peoples (Martiniano et al. 2016).

### 2.7 Elevation in TCC to TTC mutation rate is present in Mesolithic HGs and Neolithic farmers

The triplet TCC has seen a remarkable increase in mutation rates towards TTC in humans, first identified by (K Harris 2015). This signature has no known cause to date, and appears strongest in Europeans and weaker in South Asians. It was previously estimated to have started around 15,000 YBP, and its driver is most likely absent in present-day individuals (Kelley Harris and Pritchard 2017; Speidel et al. 2019), although there is considerable uncertainty about this estimate - for example, a recent study dates the onset to up to ~80k YBP depending on the demographic history used (DeWitt, Harris, and Harris 2020). One study previously quantified the signal in an early farmer (LBK) and Western HG (Loschbour), suggesting that both carried the signal, while the signal was missing in Ust’-Ishim, Neanderthals, and Denisovans (Mathieson and Reich 2017).

We first inferred the rate through time at which TCC mutates towards TTC in every individual built into our genealogy of moderns and ancients, after excluding singletons, and then quantified signal strength by calculating the area under the curve (AUC) of this rate (**Methods**). Among SGDP individuals, the quantified signal varies and is strongest in Southern Europeans such as Sardinians, who are known to have an increased affinity to early Neolithic farmers (Figure 6**a**, Supplementary Figure 10). Among the high-coverage ancients built into our *Relate* genealogies, we observe the signature in Mesolithic HGs, as well as in Neolithic and Bronze age samples, including the Yamnaya (Figure 6**a**), but infer it to be weaker in HGs and strongest in Neolithic farmers. The signal is absent in an Ethiopian HG, as expected, as well as in both the 45,000 year old Ust’-Ishim sample and the 34,000 year-old Sunghir3 sample (Figure 6**a**).

**Figure 6.**
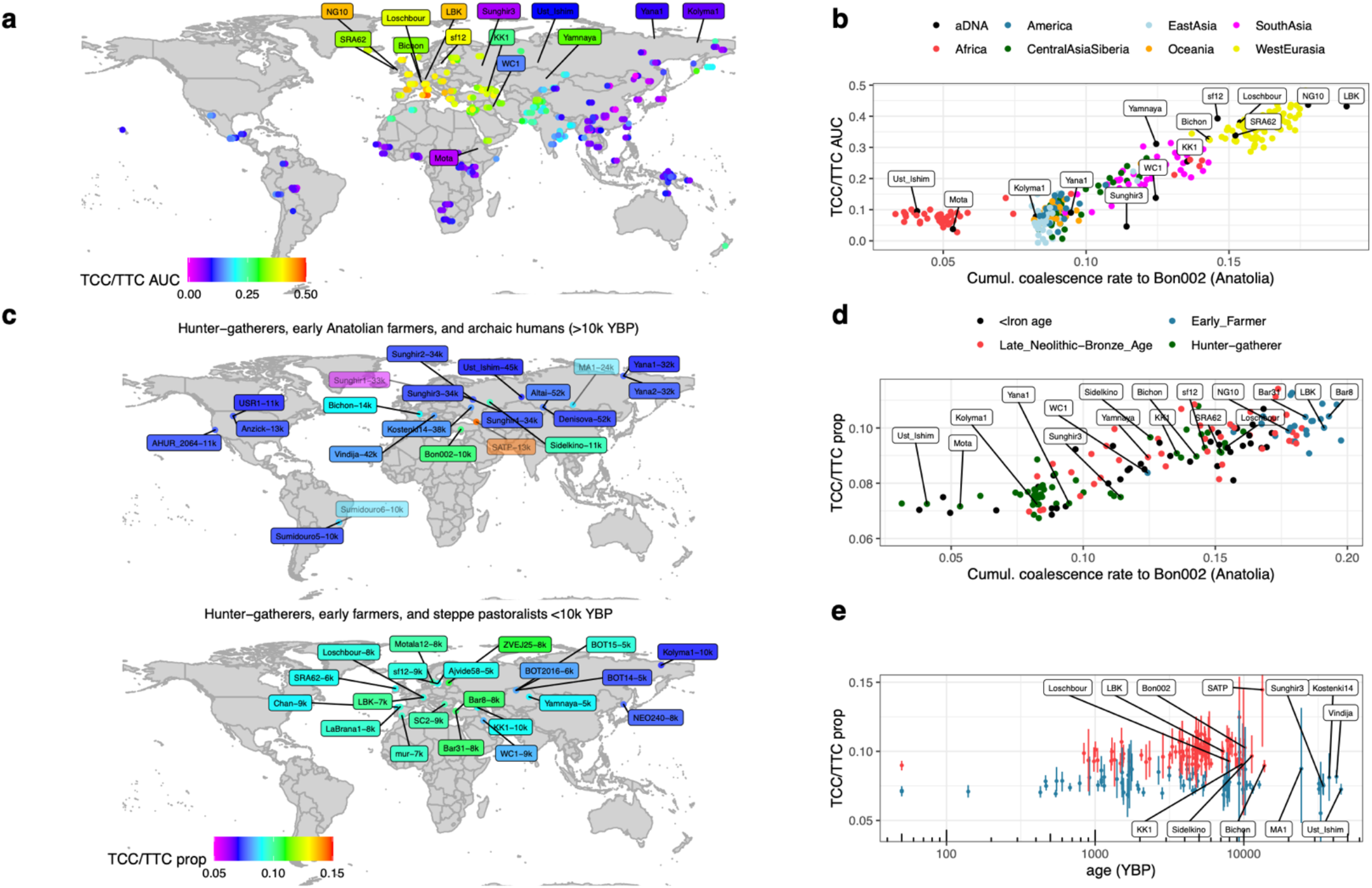
**a,** Map showing the strength of the TCC/TTC mutation rate signature, quantified by calculating the “area under the curve” (AUC) of the TCC/TTC mutation rate (**Methods**). Circles correspond to present-day individuals in the SGDP data, ancient individuals are labelled. **b)** TCC/TTC AUC plotted against the *Colate*-inferred coalescence rates to Bon002, a 10k-year-old individual from Anatolia, integrated between 14k - 50k YBP. Circles correspond to SGDP samples, labels to ancients. **c,** Map showing the TCC/TTC mutation rate signature in lower coverage ancients, quantified as the proportion of sites that are TCC/TTC relative to other C/T transitions excluding those in CpG contexts (**Methods**). Top shows a subset of samples <10k years old, bottom shows samples >10k years old (see Supplementary Figure 12 for further samples). Samples of <2x mean coverage are shown with increased transparency and number following sample ID shows sample age. **d,** Proportion of TCC/TTC sites plotted against coalescence rates to Bon002, integrated between 14k - 50k YBP. All points correspond to ancients, colour indicates their age. **e**, Proportion of TCC/TTC sites plotted against sample age. Confidence intervals are obtained using a block bootstrap. Samples are coloured using a k-means clustering (k = 2). In **c,d,e,** samples are >2x mean coverage, except for those >10k years old where we included samples >1x mean coverage.

To quantify the signal in individuals of lower coverage, we calculate the proportion of TCC/TTC mutations relative to C/T transitions in each individual, restricting to mutations ascertained in SGDP samples, of at least 4x coverage in the ancient, and dated by *Relate* to be <100k YBP (**Methods**). We confirm that signal strength is highly correlated (97%) to our AUC estimate for the high-coverage samples built into our *Relate* genealogy, where both estimates are available (Supplementary Figure 11). We do not observe the signal in Neanderthals (Prüfer et al. 2014; 2017) or Denisovans (Meyer et al. 2012), consistent with (Mathieson and Reich 2017). The signal appears already widespread in the Late Upper Paleolithic, as it is carried by Bichon, a 13,700-year-old Western HG, by Sidelkino, a 11,000-year-old Eastern HG, by SATP (Satsurblia), a 13,000 year-old Caucasus HG, and Bon002, a 10,000 year-old Anatolian Pre-Pottery individual (Figure 6**c**, Supplementary Figure 12). We note that SATP has a strong signal, however confidence intervals are large due to its lower coverage and this estimate may therefore be somewhat unreliable, although it seems clear that this individual carried the signal. The Mal’ta individual (MA1) (Raghavan et al. 2014) has a similarly large confidence interval but may not have been a carrier of this signal; WC1, a 9000-year-old Iranian farmer, who can be modelled as a mixture of a “basal Eurasian” and Mal’ta-like ancestry (Broushaki et al. 2016), and who is not closely related to Anatolian farmers, likely only carried the signal weakly, if at all. Interestingly, Chan, a 9000-year-old Iberian HG (Olalde et al. 2019) who has little ancestry related to Western HGs such as Bichon, has the weakest signal among all Mesolithic Europeans, which is at a similar level to WC1.

Already 10,000 years ago, the signal appears weaker in Western HGs compared to the Anatolian, who is among the strongest carriers of this signal (similar strength to later Neolithic individuals and present-day Sardinians) (Figure 6**e**), suggesting that the driver of this mutation rate change, which may have been of genetic or environmental nature, was already extinct by the Mesolithic. Eastern HGs have a slightly elevated signal compared to Western HGs. Moreover, the strength of the TCC/TTC signal shows a remarkable correlation with recent coalescence rates to this Anatolian individual (96% using AUC for SGDP non-Africans and 13 high-coverage ancients, 71% using TCC/TTC proportion for ancients) (Figure 6**b**, **d**), and does not correlate as well with coalescence rates to any other HG group for whom we have data (88% or 58% with Caucasus HGs (SATP), 83% or 53% with Scandinavian HGs (sf12), 76% or 37% with Eastern HGs (Sidelkino), 73% or 53% with Western HGs (Bichon), where first number uses AUC, second number uses TCC/TTC proportion) (Supplementary Figures 13). We therefore hypothesise that the signal spread through ancestors of this Anatolian individual across Europe before the arrival of farming, and subsequently arrived in Europe for a second time with Neolithic farmers.

The genetic relationship among West Eurasian HG groups in the Late Paleolithic is not fully understood and, to the best of our knowledge, current models do not include a clear source group contributing widely across these HG groups, while able to explain the strong correlation to ancestry from Anatolia. One potential source are ancestors of the Dzudzuana, a group inhabiting the Caucasus ~26k years ago (Lazaridis et al. 2018). This group is closely related to ancient Anatolians, and to a lesser extend to Caucasus HGs and may have contributed ancestry to Eastern and Scandinavian HGs before the spread of farming. The Dzudzuana have a pre-LGM common ancestor with Western HGs, including Bichon, however, placing the signal on this common ancestor lineage would not explain their signal strength difference and correlation to shared ancestry with Anatolia. Instead, one possibility is that the signal spread during the Bølling-Allerød interstadial, a brief warming following the last glacial maximum, during which Western HGs spread across Europe replacing earlier HG groups and which may have introduced gene-flow from the Near East into Europe (Fu et al. 2016).

We note that while the cause of this mutation rate elevation remains uncertain, our results would fit well with a genetic cause within a specific ancient population (for example a mutation in some repair protein, transiently present). If, alternatively, the cause is environmental, it appears highly localised in both time and place, and this seems potentially harder to explain.

## 3 Discussion

The last decade has seen an explosion in the number of sequenced ancient genomes, uncovering remarkable stories of population replacements and admixture that are associated with dramatic shifts in lifestyle arounds the world (Skoglund and Mathieson 2018). While ancient genomes are still typically available in smaller numbers and lower quality compared to genomes of present-day people, they are uniquely valuable in providing direct insight into the genetic makeup of our ancestors. We have extended the *Relate* method for inference of genome-wide genealogies to work with ancient genomes and introduced a new method, *Colate*, for inference of coalescence rates for low-coverage unphased genomes. Together, these tools enable us to harness the power of genealogy-based analyses on a wider range of samples, including those of lower quality, which were previously inaccessible.

We demonstrated, using 278 moderns of the SGDP data set, 14 high-coverage, and 430 lower-coverage ancients, that *Relate* and *Colate* can uncover dynamic population histories and evolution in the processes that drive genetic variation. The extent to which directional gene-flow occurred from groups related to ancient Anatolia into European HGs predating the spread of farming in Europe has remained controversial. We have provided two further lines of evidence that such gene-flow existed, first using coalescence rates of lineages recently coalesced between Anatolia and HGs. The TCC/TTC mutation rate elevation in all these ancient groups, and its strong correlation to inferred recent shared ancestry with Anatolia, offers complementary support that the shared ancestry detected by Colate indeed reflects recent gene exchange, given the age distribution of samples showing this mutational phenomenon.

Future avenues of research may include using genealogies for parametric inference of population histories and admixture, inspired by approaches based on site-frequency spectra (Excoffier et al. 2013; Terhorst, Kamm, and Song 2017) and F-statistics (Patterson et al. 2012; Peter 2016; Ralph, Thornton, and Kelleher 2020). Coalescence rates can be interpreted as a function of gene flow (or the lack thereof); for instance, (Wang et al. 2020) have recently developed a method that infers migration rates through time given pairwise coalescence rate estimates. Genealogies of modern individuals have proven to be very powerful in quantifying positive selection (Speidel et al. 2019; Stern, Wilton, and Nielsen 2019; Stern et al. 2021) and genealogies including ancient genomes should further boost power.

While *Colate* has made it possible to leverage genealogies for the study of low-coverage genomes possible, we ideally would like to incorporate such genomes directly into genealogical trees. This is currently not possible, however recent work building on the tsinfer methodology (Kelleher et al. 2019) provides an alternative approach that constrains the age of ancestral haplotypes using low-coverage ancient genomes to infer genome-wide genealogies for phased sequences (incl. ancients and moderns) (Wohns et al. 2021). A possibility for making lower coverage ancient genomes, or indeed hybrid capture array data, accessible to these methods is imputation (Rubinacci et al. 2020; Hui et al. 2020). A potential concern is that imputation may introduce biases, particularly in ancient genomes with ancestries that are not well reflected in modern groups. These biases are often difficult to assess. Because *Colate* does not require imputation, we expect that it will be a useful tool to investigate such biases in future.

## 4 Methods

### 4.1 Colate

Coalescence rates are inferred by attempting to maximise the following likelihood using an expectation-maximisation (EM) algorithm. For any derived mutation carried by a reference chromosome *j*, we ask whether this mutation is shared by the target chromosome *i* which we denote by an indicator variable *I_ℓ,ij_*(*ℓ* indexing SNPs). We multiply across SNPs, such that no phase information is required to compute the likelihood. To obtain coalescence rates between groups of individuals, we also multiply the likelihood across homologous chromosomes in both the target and reference groups. To calculate within-individual coalescence rates, the method assigns one allele to each category, at random at every SNP. When input is specified in BAM format (as reference-aligned reads), we multiply across reads. The maximum likelihood estimate is then given by 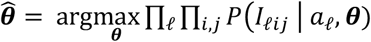, where ***θ*** denotes piecewise-constant coalescence rates and *a_ℓ_* is the age of the *ℓ*th mutation, which we assume to be known here, but have to integrate out in practice.

To integrate out mutation age, we assume neutrality of every mutation, implying that its age is uniformly distributed on the branch onto which it maps. The EM algorithm requires us to integrate out mutation age conditional on sharing or not sharing between target and reference chromosomes. This theoretically implies a deviation from the uniform distribution. This deviation is strongest for mutations that are singletons in the genealogy used to date these mutations and are shared between sequences in the target and reference chromosome sets; in this case, knowledge of sharing implies that the mutation is older than the coalescence time of these chromosomes, biasing mutation age upwards compared to a uniform distribution (Supplementary Figure 14). We use an empirical approach to sample mutation ages for these shared singletons and use the uniform distribution for all other mutations in practise, which we demonstrate is a reasonable approximation (SI). Moreover, we note that the *Colate* approach requires the inclusion of sites fixed and derived in all samples used for inferring the genealogy, as samples can, in theory, coalesce into the root branch. To obtain an approximate upper bound on the age of such mutations, we fix the time to the most recent common ancestor (TMRCA) to an outgroup (10M YBP for human-chimpanzee in this study).

We bin mutation ages into a discrete time grid to reduce computation time of the EM algorithm. As a result, the algorithm only requires the number of shared and not-shared mutations in each time grid as input; compilation of this input data is linear in sample size and number of mutations. Once in this form, the input data to the EM algorithm, and hence the computation time of the EM algorithm, is independent of sample size or the number of mutations and takes approximately 5 seconds (~40 seconds including parsing of the data).

### 4.2 Simulations

We used stdpopsim to simulate genomes with different demographic histories (Adrion et al. 2020) and hotspot recombination rates to evaluate *Relate* and *Colate*. For *Colate*, we additionally require an outgroup to determine mutations that are fixed in all samples. Instead of simulating an outgroup explicitly, we fixed the time to the most recent common ancestor (TMRCA) *t_out_* to the outgroup (*t_out_* = 10M years in our simulations), and sampled the number of fixed mutations in any given region as a Poisson distributed random variable with mean *μl*(*t_out_* – *t_sample_*), where *μ* is the per base per generation mutation rate, *t_sample_* is the TMRCA of the sample in this region and *I* is the number of base-pairs in this region. If *t_sample_* was greater than *t_out_*, we sampled no fixed mutations. We then chose the base-pair positions of these fixed mutations uniformly at random with replacement within the corresponding region. For simplicity, we assumed a two-state mutation model, such that a repeat mutation at one genomic site return to the original state.

Supplementary Figure 2 shows the performance on a zigzag history (Schiffels and Durbin 2014), demonstrating near perfect recovery of coalescence rates when using true mutation ages in *Colate*, and high accuracy when mutation ages are sampled given a genealogy; the discrepancy highlights that our sampling distribution of mutation age given a genealogy (**Methods**, Supplementary Information) is reasonable but not exact.

We also simulated the multi-population model of ancient Eurasia from the stdpopsim package, which was fitted using real human genomes (Kamm et al. 2020). We simulated 200 haploid sequences in each of three modern human groups (Mbuti, Sardinian, Han), as well as four ancient Eurasians (LBK, Loschbour, Ust’-Ishim, MA1) and a Neanderthal (two haploid sequences in each group) (Figure 2**a**). From this simulation, we obtained true genealogical trees and *Relate* trees for all samples. In addition, we inferred a separate set of *Relate* trees using only the three modern human groups (Mbuti, Sardinian, Han), which we used to date mutations for *Colate*.

*Colate* recovered within and across group coalescence rates accurately compared to the corresponding direct MLEs calculated on true or Relate-inferred trees (Figure 2**a**). In particular, these coalescence rates clearly captured the admixture from Neanderthals into an ancestral Eurasian lineage, as well as more recent genetic structure, such as separation of the Loschbour HG and early farmer lineages, represented by LBK. We observed a closer affinity of the Loschbour HG to modern-day Sardinians, compared to LBK, consistent with modern Sardinians being an admixture of HG and farmer ancestry in this simulation.

One case for which *Colate* performed less well compared to direct MLEs obtained from *Relate* trees is the crosscoalescence rates between Neanderthals and Mbuti, calculated by assigning the Neanderthal as reference and Mbuti as target. This is because the genealogy used to date mutations contains only variants segregating in the three modern groups and therefore captured almost none of the Neanderthals variation that postdates the Neanderthal-Mbuti split. In this case, it would therefore be preferable to instead assign Mbuti as reference.

### 4.3 Evaluating *Colate* on downsampled high-coverage genomes

We evaluated the performance of *Colate* on low-coverage sequencing data, by comparing estimates obtained from downsampled BAM files (Figure 2**b**). To date mutations, we constructed a genealogy containing 25 diploid samples from each of the three 1000 Genomes populations - YRI (Yobura in Ibadan, Nigeria), CEU (Northern and Central European ancestry individuals from Utah, USA), and CHB (Han Chinese from Beijing, China) (The 1000 Genomes Project Consortium 2015), downloaded from http://ftp.1000genomes.ebi.ac.uk/vol1/ftp/release/20130502/. We then chose four 1000 Genomes samples that do not overlap the genealogy as target chromosomes (HG00096, HG00268, NA18525, NA19017) and included the remaining samples in groups YRI, CEU and CHB in the reference chromosomes set. The BAM files of these four genomes were obtained from ftp://ftp.1000genomes.ebi.ac.uk/vol1/ftp/technical/working/20140203_broad_high_cov_pcr_free_validation/ma_tching_LC_samples_bwamem/ subsequently downsampled using SAMtools v1.9 (H. Li et al. 2009).

Across a wide range of mean coverages, *Colate*-inferred coalescence rates remained unchanged and nearly identical to rates inferred using called genotypes (VCF). To obtain 95% confidence intervals, we used a block bootstrap, dividing the genome into 20Mb blocks, and resampling 100 times. Confidence intervals become wider for lower coverage sequencing data; encouragingly, we could infer meaningful coalescence rates between a target sequence of 0.01x mean coverage and the reference VCFs.

We additionally evaluated *Colate* when both target and reference samples are of low coverage by calculating the coalescence rates between LBK, a 7200 year old early European farmer, and Loschbour, a nearly 8000 year old Mesolithic Western HG (both >14x coverage) (Lazaridis et al. 2014) using a genealogy for SGDP to date mutations. We downsampled both individuals to a minimum of 0.1x mean coverage (Supplementary Figure 3). While inference of coalescence rates became challenging when both genomes are at 0.1x, estimates still appeared reasonably accurate and unbiased.

### 4.4 Data

#### 4.4.1 Simons Genome Diversity Project Data

We downloaded phased haplotypes for 278 individuals from https://sharehost.hms.harvard.edu/genetics/reich_lab/sgdp/phased_data/PS2_multisample_public/, and rephased these jointly with high coverage ancients (Section 4.4.2) using SHAPEIT4 (Delaneau et al. 2019). We first used the 1000 Genomes Project (1000GP) reference panel (http://ftp.1000genomes.ebi.ac.uk/vol1/ftp/release/20130502/) to phase all sites overlapping with 1000GP and then internally phased all remaining sites, while keeping the already phased sites fixed.

#### 4.4.2 Ancient genomes data

We downloaded 430 ancient genomes for use in this study (Supplementary Table 1). All samples had a genomewide mean coverage of 0.5x or more. We selected 14 high coverage ancient genomes (mean genomic coverage > 7.8X) for *Relate* analysis.

For the 14 high coverage genomes (Supplementary Table 1) genotypes were called using samtools mpileup (input options: -C 50, -Q 20 and -q 20) and bcftools call --consensus-caller with indels ignored (H. Li 2011). A modified version of the bamCaller.py script from https://github.com/stschiff/msmc-tools was used to output variant sites. We generated a mask for each ancient genome, declaring only sites with at least 5X coverage and below twice the mean genomic coverage as passing.

We merged these 14 ancient genomes with the 278 Simon Genome Diversity Project samples to infer joint genealogies using *Relate.* We applied a conservative mask, declaring only sites passing in all of the 14 ancients, as well as a universal mask file provided with the SGDP data set, as passing. The SGDP universal mask was obtained from https://reichdata.hms.harvard.edu/pub/datasets/sgdp/filters/all_samples/.

### 4.5 Joint genealogies of ancients and moderns

We inferred joint genealogies of ancients and moderns using our updated *Relate* algorithm (Supplementary Information). We used all mutations, excluding those in CpG contexts, to infer tree topologies and restricted to transversion only for inference of branch lengths. We therefore used a reduced mutation rate of 3e-9 per base per generation. We used a recombination map obtained from https://mathgen.stats.ox.ac.uk/impute/1000GP_Phase3.html and realigned alleles relative to an ancestral genome obtained from ftp://ftp.1000genomes.ebi.ac.uk/vol1/ftp/phase1/analysis_results/supporting/ancestral_alignments/. We used default parameters in *Relate* otherwise.

To infer branch lengths, we used a precomputed average coalescence rate estimate obtained by applying *Relate* to the 278 SGDP moderns. To compute these coalescence rates, we jointly sampled branch lengths and effective population sizes using our updated iterative algorithm, which we show can be interpreted as an approximate EM algorithm for finding maximum likelihood coalescence rates. This approximate EM algorithm samples genealogies using *Relate* instead of integrating over all possible genealogies (see Supplementary Information Section B). To obtain a coalescence rate estimate that matches the mutation rate used for inferring the genealogy of ancients and moderns, we inferred branch lengths using transversions only and set the mutation rate to 3e-9 per base per generation.

### 4.6 *Colate*-inferred coalescence rates for SGDP and 430 ancients

We inferred coalescence rates for pairs of ancient individuals using *Colate*, restricting to transversions only. For each pair of samples, when given as a VCF file, we applied the respective mask files. When a sample was given in BAM file format, we accepted a read whenever mapping quality exceeded 30, read length exceeded 34 bps, and there were fewer than three mismatching sites. We further excluded 2 base-pairs at each end of a read and restricted our analysis to sites where at most two different alleles were observed.

To date mutations, we used a *Relate*-inferred genealogy of the SGDP dataset. As the degree of inbreeding varied across SGDP individuals (main text) and to avoid biases in mutation ages resulting from extensive inbreeding in some individuals, we selected one haploid sequence from each individual. We jointly fitted branch lengths and coalescence rates using a mutation rate of 1.25e-8 per base per generation.

### 4.7 Calculation of mutation rate

We calculated mutation rates for 7 6 mutation triplets in each individual, after excluding any singletons and terminal branches in our genealogy. We only considered mutation triplets that are not in a CpG context, which excludes 20 of 96 possible triplets. To remove trends shared across mutation triplets, we divided the TCC/TTC mutation rate by the average over all triplets (excl. CpG contexts) in each epoch, to obtain the mutation rate relative to the average mutation rate.

To calculate the area under the curve for the TCC/TTC mutation rate signature, we first scaled the mutation rate in each individual by the average over the time interval [1e5,1e6] YBP. We then calculated the area under the curve between 14k to 1M years BP. For samples that are older than 14k years (Ust’-Ishim, Sunghir3, and Yana1), we extrapolated the earliest value to 14k YBP. We then subtracted the equivalent value of a constant mutation rate from this AUC, such that any sample without the elevation in TCC/TTC mutation rates is expected to have an AUC of 0.

### 4.8 Quantifying the TCC/TTC signal in lower coverage individuals

We quantified the TCC/TTC signal in lower coverage individuals (>2x mean coverage) by restricting to sites segregating in our SGDP genealogy that we also used to date mutation in *Colate.* We additionally restricted to sites where the age of the upper coalescence event of the branch onto which the mutation maps is <100k YBP. For each sample, at any such site, we then further restricted to sites where at least four mapping reads, and added a count towards a mutation category if at least four reads supported the derived allele. In this way, we counted the number of sites that are likely to be in heterozygous or homozygous state for the derived allele. We finally calculated the proportion of such sites, relative to any C/T transitions, excluding those in CpG context. We calculated confidence intervals using a block bootstrap with block size of 10Mb.

### 4.9 Calculation of pairwise F2 statistics

We calculated F2 statistics between ancients for comparisons to matrices of pairwise coalescence rates (used in Supplementary Figure 6). To calculate F2 statistics, we first made pseudohaploid calls for each individual using “pileupcaller” (https://github.com/stschiff/sequenceTools), where we restricted to 1240k ascertained genomic sites known to be varying among present-day humans (Mathieson et al. 2015). We then merged individuals using “mergeit” (https://github.com/DReichLab/EIG). To calculate F2 statistics, we used the R package admixtools2 (https://github.com/uqrmaie1/admixtools).

## Supporting information

SITable1

## Acknowledgements

We thank David Reich, Richard Durbin, Iain Mathieson, Mateja Hajdinjak, Anders Bergström, Simone Rubinacci, and Clare E. West for helpful comments, discussions, and ideas. L.S. acknowledges support provided by the Sir Henry Wellcome fellowship [220457/Z/20/Z]. L.S., G.H. and S.R.M acknowledge funding from the Wellcome Trust [200186/Z/15/Z] and S.R.M. acknowledges support provided by the Wellcome Trust Investigator Award [212284/Z/18/Z]. P.S. is supported by the Francis Crick Institute core funding (FC001595) from Cancer Research UK, the UK Medical Research Council, and the Wellcome Trust. P.S. is also supported by the European Research Council (grant no. 852558), a Wellcome Trust Investigator award (217223/Z/19/Z) and the Vallee Foundation. This research was funded in whole, or in part, by the Wellcome Trust. For the purpose of Open Access, the author has applied a CC BY public copyright licence to any Author Accepted Manuscript version arising from this submission.

## Software availability

Relate: https://myersgroup.github.io/relate/

Colate: https://github.com/leospeidel/Colate

## Supplementary Information

### A. Colate

#### A.1 Notation

We will use the following notation in this section:

- *e* indexes epochs
- *τ_e_* denotes the lower boundary of epoch *e*
- *θ*(*t*) denotes coalescence rates through time and takes the form 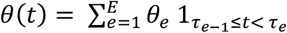, where 1 is the indicator function. We write ***θ*** = (*θ_e_*)_*e*=1:,…,E_.
- *I_ℓ_* is the indicator of sharing/non-sharing of a mutation at site *ℓ*
- *t_ℓ_* is the coalescence time at SNP *t*, which is unknown
- *a_ℓ_* is the age of a mutation at site **ℓ** and *l_ℓ_* and *u_ℓ_* are the lower and upper times of the branch onto which this mutation maps.

#### A.2 Overview of the Colate method

Throughout, we assume to have one reference and one target sequence. In cases where we have multiple reference or target sequences (e.g., in non-haploid organisms, or groups of individuals), we use a composite likelihood approach and multiply the likelihood across individuals. Colate can be applied to reference-aligned read data directly by constructing a composite likelihood that multiplies over reads. We also use a composite likelihood approach across genomic sites and therefore require no phase information for non-haploid organisms.

Epoch boundaries *τ_e_* are prespecified parameters and we assume that the coalescence rate is given by a piecewiseconstant function 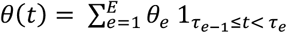. We aim to find a maximum likelihood estimate of the coalescence rates ***θ*** = (*θ_e_*)_*e*=1,…,*E*_.

For any mutation carried by the reference sequence, we observe whether the mutation is also carried by the target sequence. This is our observed data and is stored in the indicator variable *1_{_* equaling 1 if mutation *ℓ* is shared and 0 if it is not shared. In the following Expectation-maximisation (EM) algorithm, the coalescence time *t_ℓ_* at SNP *ℓ* between the target and the reference sequence is the unobserved latent variable which we will integrate out. In the first part, we will assume that mutation age *a_ℓ_* is known and we will extend our method to the case when mutation age is unknown in the second part. The EM algorithm maximises Π_ℓ_ *P*(*I_ℓ_* | *a_ℓ_ **θ***) (*ℓ* indexing SNPs) with respect to coalescence rates ***θ***, outputting an approximate maximum likelihood estimate (MLE) 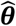. We obtain uncertainty estimates around this MLE using a block bootstrap on genomic regions.

#### A.3 Expectation-Maximisation algorithm with known mutation ages and genotypes

We assume that mutation ages *a_ℓ_* are known. Then, the loglikelihood of ***θ*** given the data *I_ℓ_* and latent variable *t_ℓ_* is

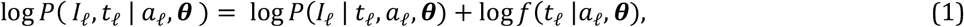

where *P*(*I_ℓ_* | *t_ℓ_*, *a_ℓ_, **θ***) is a step function given by

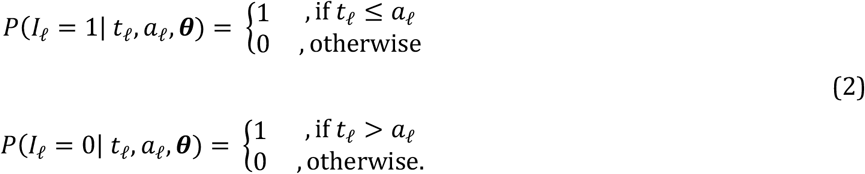

This step function reflects our infinite-sites assumption: a mutation can only be shared if it is older than the time to the most recent common ancestor (TMRCA) between the target and reference sequence and it can only be not shared if it is younger than the TMRCA. In particular, this step function does not depend on ***θ***. The density of coalescence rates ***θ*** is a non-homogeneous exponential given by the standard coalescent, which does not depend on mutation age *a_ℓ_*; it is given by

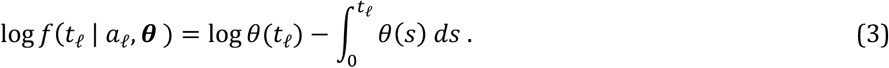

By using that 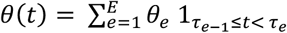, we can rewrite Eq. (3) as

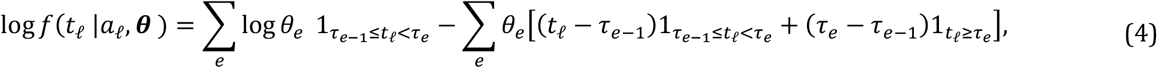

where 1_*x*_ denotes the indicator function equaling one if and only if *X* is true and 0 otherwise.

The EM-algorithm requires us to integrate out the latent variable *t_ℓ_* conditional on the data and the coalescence rates of the previous iteration, denoted by ***θ***^(***k***)^. Substituting Eq. (4) in Eq. (1) and taking the expectation, we obtain

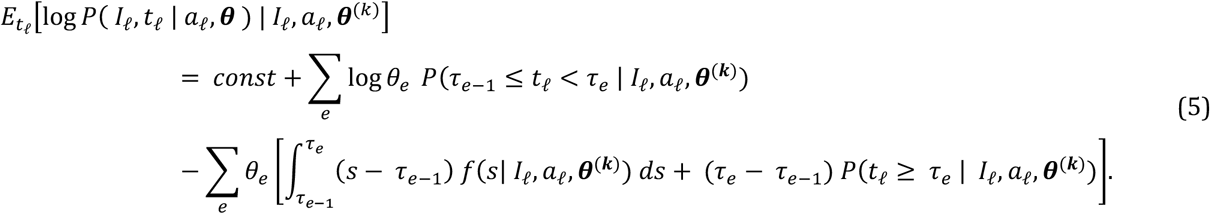

Equation (5) is the expected log-likelhood for one SNP. We use a composite likelihood across SNPs, such that the expected log-likelihood genome-wide is a sum of Eq. (5) across all SNPs. To complete the EM update, we maximise the expected loglikelihood with respect to ***θ*** to obtain our updated estimate ***θ***^(***k***+1)^ By finding the root of the first derivative with respect to *θ_e_*, we obtain

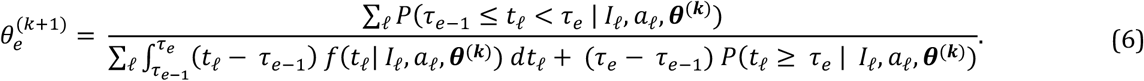

The numerator of Eq. (6) is the probability that the coalescence event occured in epoch *e*. The denominator of Eq. (6) is the opportunity (or expected branch length) of the coalescence event happening in epoch *e*. Evaluation of Eq. (6) requires calculating integrals of 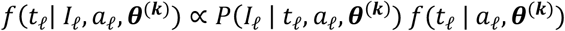, which is given by Eqs. (1)(3) and is effectively an integral of the exponential prior density of coalescence times over an adjusted domain that excludes coalescence events incompatible with the data (i.e., sharing/non-sharing of the mutation).

In practice, we use a discrete time grid to calculate Eq. (6). By doing so, we can bin SNPs by age bins, such that we only have to calculate a constant number of integrals (not growing with the number of SNPs) to evaluate Eq. (6).

When multiple target and/or reference sequences are used, we precompute the how often the mutation is shared and non-shared by age bin. Given these precomputed values, evaluation of Eq. (6) is not dependent on the number of SNPs or the number of target and reference sequences. Counting how often a mutation is shared and non-shared only requires computing the derived allele frequencies in the target and reference sample and is given by 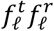 and 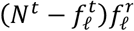, respectively, where 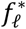 denotes the derived allele frequency and *N*\* the number of sequences.

Overall, the computational complexity of this EM algorithm is constant with respect to number of SNPs and number of sequences, beyond calculating the number of shared/non-shared mutations by age bin, which itself takes linear time (in number of SNPs and number of sequences) and requires little computation beyond parsing the data and computing derived allele frequencies.

#### A.4 Expectation-Maximisation algorithm with unknown mutation ages and known genotypes

In practice, mutation ages are unknown and we infer mutation ages using a genealogy. This genealogy is inferred for individuals that are usually distinct from the reference sequences in the EM algorithm, e.g., in practice, we might use a large sample to infer a genealogy to date mutations, and subsequently infer coalescence rates between targets and a subset of the sequences used to infer the genealogy, or two target sequences. A genealogy will limit the mutation age *a_ℓ_* to a range between the lower and upper boundaries of the branch onto which the mutation maps, which we denote by *l_ℓ_* and *u_ℓ_*. We modify our EM algorithm and treat mutation age as an additional latent variable, in addition to the coalescence time *t_ℓ_*, such that Eq. (1) is updated to

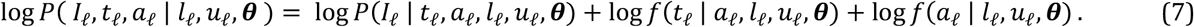

Here, *P*(*I_ℓ_* | *t_ℓ_,a_ℓ_,l_ℓ_,u_ℓ_,**θ***) is still the same step function and does not depend on *l_ℓ_*, *u_ℓ_*, and ***θ***. The density of mutation ages *f*(*a_ℓ_* | *l_ℓ_, u_ℓ_ **θ***) is given by the uniform distribution between *l_ℓ_* and *u_ℓ_* and does not depend on ***θ***. We note that *f*(*t_ℓ_* | *a_ℓ_, l_ℓ_, u_ℓ_ **θ***) is no longer given by a non-homogeneous exponential, as we are conditioning on *l_ℓ_* and *u_ℓ_*.

Using Eq. (7), the expected log-likelihood is given by

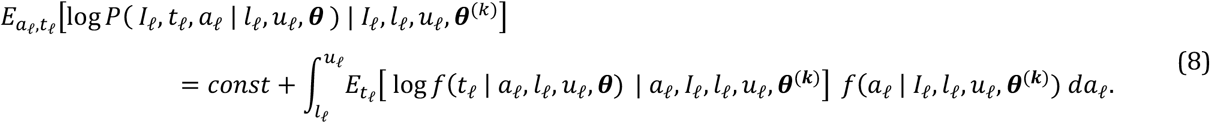

Instead of evaluating the integral over *a_ℓ_*, we will attempt to sample *a_ℓ_* from the distribution *f*(*a_ℓ_* | *l_ℓ_, l_ℓ_, u_ℓ_, **θ***^(*k*)^)*da_ℓ_*.

If we can sample *a_ℓ_* in an unbiased way, we on average “know” the age of the mutation and expect

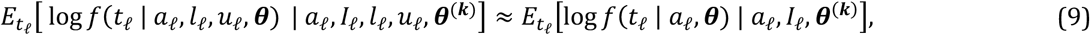

which will bring us back to the case where mutation age is known.

#### A.5 Sampling mutation ages given genealogical constraints

It is key to sample from *f*(*a_ℓ_* | *l_ℓ_, l_ℓ_, u_ℓ_, **θ***^(***k***)^) in an unbiased way. Here we illustrate an approximate approach that works well in practice. We use Bayes’ theorem and obtain

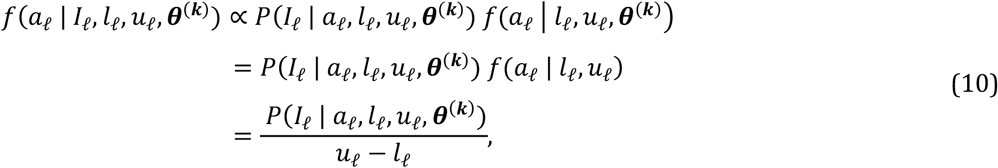

where we use that unconditionally, the age of a mutation is uniformly distributed between *l_ℓ_* and *u_ℓ_*. We are therefore interested in the functional form of *P(I_ℓ_* | *a_ℓ_, l_ℓ_, u_ℓ_, θ^(k)^).* At first glance, it seems as if we can approximate

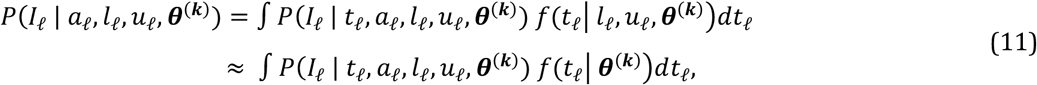

where *P*(*I_ℓ_ | t_ℓ_,a_ℓ_,l_ℓ_,u_ℓ_,**θ***^(***k***)^) is the step function taking 1 if sharing (or non-sharing) is compatible with coalescence time and mutation age (defined in Eq. (2)) and the approximation is based on *f*(*t_ℓ_*| *l_ℓ_,u_ℓ_,**θ***^(***k***)^) ≈ *f*(*t_ℓ_*| ***θ***^(***k***)^), with the latter being the coalescent prior. However, this approximation introduces a bias, which will invalidate our earlier approximation in Eq. (9).

Instead, we argue that

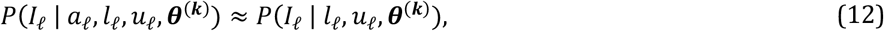

where the right-hand side does not depend on mutation age *a_ℓ_*, implying that Eq. (10) is the uniform distribution on [*l_ℓ_, u_ℓ_*). Intuitively, this means that the probability of sharing (or non-sharing) does not depend on the mutation age, beyond conditioning on boundaries of the branch it falls on; this should be accurate if the probability of coalescing into this branch is negligible, and the more likely scenario is that coalescences happen either before *l_ℓ_* or after *u_ℓ_*. We show that empirically, this is the case in Supplementary Figure 14.

As Supplementary Figure 14 shows, approximating *f*(*a_ℓ_* | *l_ℓ_, l_ℓ_,u_ℓ_,**θ***^(***k***)^) by the uniform distribution is reasonable in most cases. An important exception are shared singletons. The age of a shared singleton is not well approximated by a uniform distribution, because the target and reference sequence coalescence into the branch onto which this singleton maps with certainty.

We therefore treat shared singletons separately by sampling from the following empirical distribution of singleton age. For shared singletons, we therefore approximate the distribution function of its age *a* by

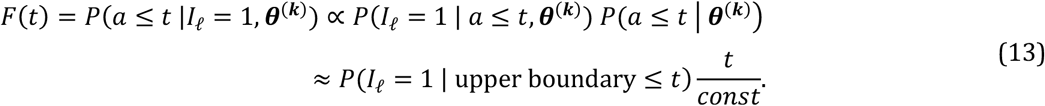

We calculate *P*(*I_ℓ_* — 1 | upper boundary ≤ *t*) empirically using the fraction of shared singletons with upper boundary not greater than t. The term *t/const* assumes that a mutation happens sometime between time 0 and the time to the shared ancestor with an outgroup, such that unconditionally of sharing/non-sharing, the distribution of the age of a singleton is approximately uniform. Using Eq. (13), we can now sample the age of a singleton conditional on whether it is shared, using the inverse-transform trick, such that *a ~ F*^-1^(*U*), with *U* being a uniform random variable on [0,1].

### B. Relate: Approximate EM algorithm for inferring coalescence rates from a genealogy

In (Speidel et al. 2019), we described an iterative algorithm for estimating branch lengths and coalescence rates; this algorithm iteratively inferred maximum likelihood coalescence rates given a tree, and then used these coalescence rates to reestimate branch lengths. This algorithm worked well in practise, but was heuristic.

Here, we describe how a minor modification of this algorithm can be interpreted as an approximate EM algorithm that attempts to find the maximum likelihood coalescence rates for given data, essentially integrating out the possible genealogical histories by sampling these using Relate.

As before, we let ***θ*** = (*θ_e_*)_*e*=1,…,*E*_ be the coalescence rates in epochs *e* = 1, …, *E*. Here, we describe a method that is slightly modified from the method in Speidel et al. (2019) for inferring coalescence rates using genealogies. We aim to find the maximum likelihood estimate

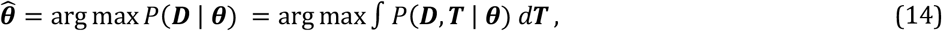

where ***D*** is the observed genetic variation data and ***T*** = (*T_ℓ_*)_*ℓ*_ is the collection of local genealogies, which we treat as unobserved latent variables in the following EM algorithm. For one marginal tree *T_ℓ_* the log likelihood is given by

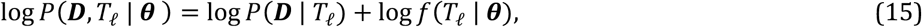

where *P*(***D*** |*T_ℓ_*) is typically given by a Poisson model (mutations happening at a constant rate *μ*), which does not depend on coalescence rates ***θ***, and *f*(*T_ℓ_* | ***θ***) is the coalescent prior of the marginal tree given coalescence rates. Denoting our estimate of the coalescence rate in step *k* of the EM algorithm by ***θ***^(***k***)^ and multiplying likelihoods across trees, the update of the EM algorithm is given by

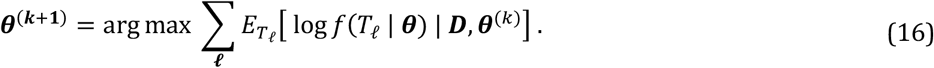

Integrating formally over marginal trees given the data is difficult, so instead we use genealogies sampled by Relate. In this approach, tree topology is fixed, and branch lengths are sampled from the posterior distribution given the data (mutations mapped to branches). In Speidel et al. (2019), where the algorithm for estimating coalescence rates was formulated in a more heuristic way, we instead used posterior mean branch lengths; by sampling branch lengths the algorithm is now an approximate EM algorithm.

Another difference to Speidel et al. (2019) is that we use the full coalescent prior in our approach here, whereas we used the coalescent prior for two haploid sequences in Speidel et al. (2019) and then averaged over all pairs of haploid sequences afterwards. Denoting by *t_f_* the time of the coalescence event reducing the number of lineages from *j* + 1 to *j* back in time, the coalescent prior is given by

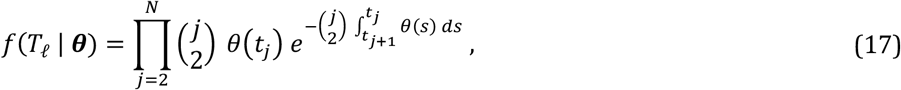

where coalescence rates are piecewise constant, i.e., 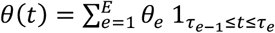. Applying this to the logarithm of Eq. (17), we obtain

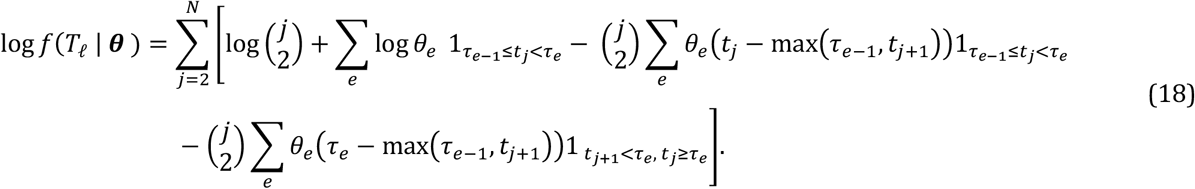

Substituting Eq. (18) in Eq. (16) and assuming that we only sample one marginal tree per locus (which is the case in practice), we obtain

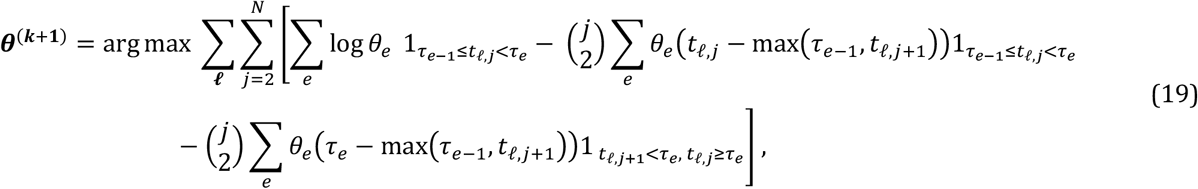

where *t_ℓ,j_* denotes the coalescence time of the event reducing the number of lineages from *j* + 1 to *j* in the *ℓ*th tree. Calculating the root of the first derivative with respect to *θ_e_* gives

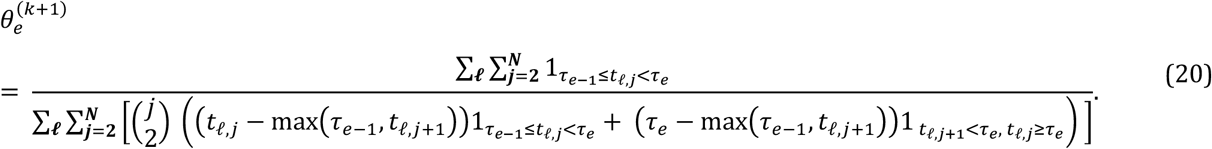

Similarly to Eq. (6), the numerator of Eq. (20) counts the number of coalescence events happening in epoch *e* and the denominator of Eq. (20) measures the total opportunity for a coalescence event in this epoch. We note that if *N* = 2, *t*_*ℓ*,2_ is the coalescence time between two sequences and *t*_*ℓ*,3_ — 0, such that Eq. (20) reduces to the estimator derived in Speidel et al. (2019) for two haploid sequences.

### C. Adapting Relate to build genealogies including ancient genomes

We modified the tree builder for constructing tree topologies and the branch length sampling scheme in our Relate method; the remainder of the method is unchanged and we refer the reader to Ref. (Speidel et al. 2019) for details of the method.

#### Tree builder for ancient genomes

The challenge with including ancient genomes is that these impose hard constraints on branch lengths and coalescence times; any coalescence events has a minimum age which is the maximum age of its descendants. We therefore modified the tree builder to discourage coalescences between contemporary and non-contemporary genomes when there is no strong evidence for this coalescence.

To do this, we calculate a preliminary date for coalescence events while inferring tree topology. Our tree builder constructs local genealogical trees bottom-up and we assign an age by calculating the expected time under the coalescent model, given the number of remaining lineages and a pre-specified effect population size as input. We then only allow coalescences between non-contemporary samples and other lineages, if the age of that lineage exceeds the sampling age of the non-contemporary sample, except when this is the only feasible coalescence event.

Identification of feasible coalescence events is identical to before, where we find pairs of lineages that are mutually minimal in a non-symmetric distance matrix calculated using a modified chromosome painting hidden Markov model (Speidel et al. 2019; N. Li and Stephens 2003). Whenever we have more than one feasible pair of lineages (that are allowed to coalesce in our rule for non-contemporary samples above), we choose the pair with minimal distance in the symmetrised distance matrix.

#### Markov-chain Monte Carlo sampler for branch lengths

We modified the MCMC update rules to allow for non-contemporary samples. This MCMC algorithm samples from the following posterior distribution

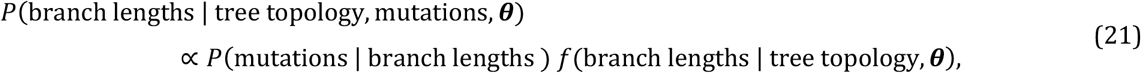

where *P* (mutations | branch lengths) is the likelihood function given by a Poisson model with a constant mutation rate and *f*(branch lengths | tree topology, *θ*) is the coalescent prior on branch lengths.

We have two ways of proposing new branch lengths, which are chosen at random with probability 0.4 and 0.6, respectively.

#### Swapping the times of two events (same as before)

This step is unchanged. We choose two events at random and propose a switch of their coalescence times, if this does not violate tree topology. The times while *k* lineages remain are unchanged and the update step only requires recalculation of the likelihood function of the six branches that have been proposed to change in length (two daughter and one parent branch for each of the two events).

#### Update a single event between older daughter and parent (new)

For modern samples, we additionally used an update step that proposed a new time for the time while *k* ancestors remain. Here, we replace this step with a new update step that proposes to only change the timing of one coalescence event to anywhere between its older daughter event and parent event. We first choose one coalescence event at random. Defining the age of the older daughter coalescence event by *t*_d_ and the age of the parent coalescence event by *t*_p_, the proposed age of the chosen event is drawn from a uniform distribution on [*t*_d_, *t*_p_).

The acceptance probability in a Metropolis-Hastings type MCMC sampler is given by the ratio of proposal probabilities of the old and new age of the chosen coalescence event, multiplied by the ratio of posterior probabilities of the old and new branch lengths. Conveniently, the proposal distribution is symmetric with respect to old and new ages, such that the ratio for the proposal probabilities is 1. It remains to evaluate the ratio of posterior probabilities of branch lengths, which are given by Eq. (21).

## Supplementary Figures

**Supplementary Figure 1.**
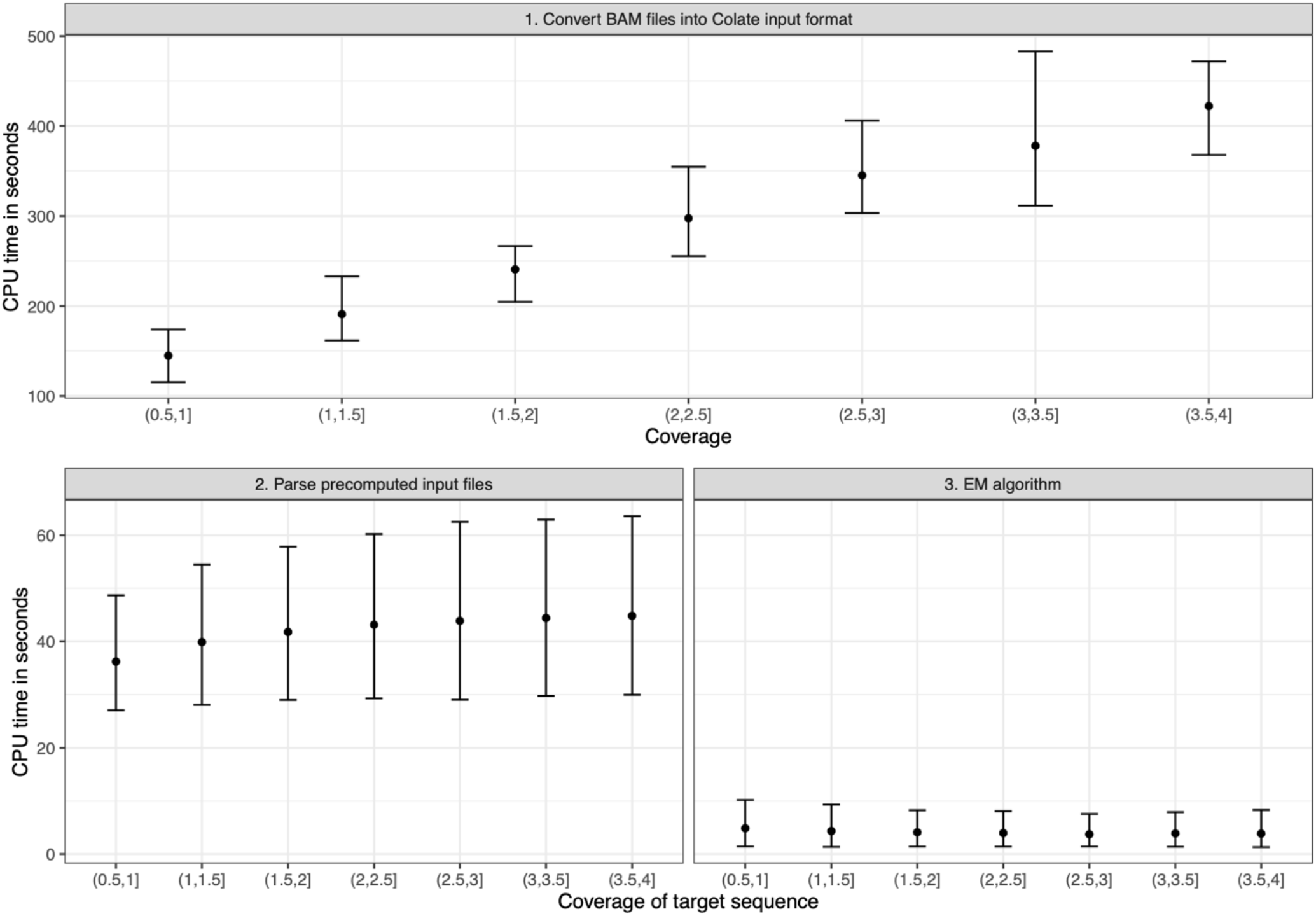
Runtime of Colate on ancient genomes of <4x coverage, using mutations dated in a genealogy estimated using SGDP individuals (**Methods**). Step 1 converts BAM files into an input file format used for Colate, that stores the number of reads supported each allele at sites dated in the genealogy. This step is linear in coverage. Step 2 parses two sequences that were each processed using Step 1 and scales linearly with the number of mutations used in the analysis, which scales somewhat with increasing coverage as more mutations are included in the analysis. Step 3 infers maximum likelihood coalescence rates using an EM algorithm that is now independent of input sequence coverage and the number of mutations used in the analysis. The x-axis in steps 2 and 3 denotes the coverage of the target sequence; coverage of the reference sequence ranges from 0.5x to 4x and is reflected in the error bars.

**Supplementary Figure 2.**
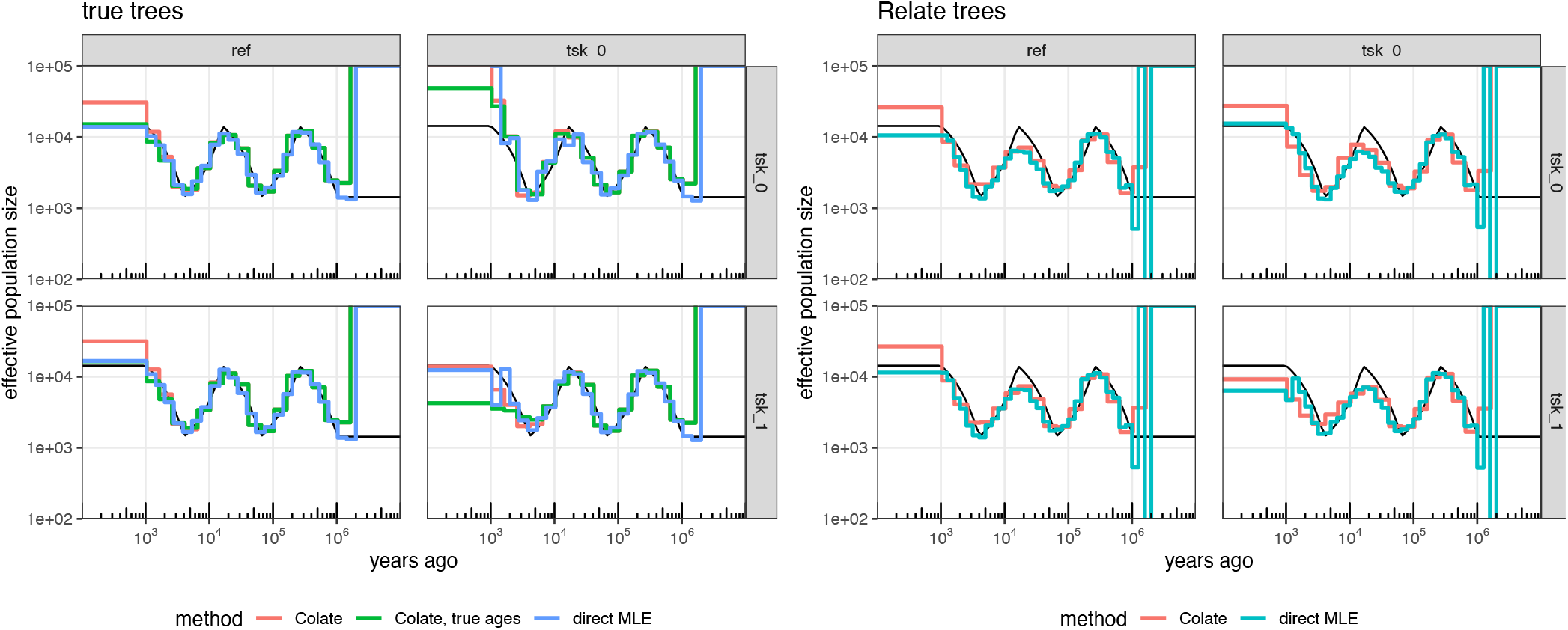
Inferred effective population sizes using (**a**) true trees and (**b**) Relate trees for a stdpopsim simulation equivalent to whole human genomes of 102 diploid sequences with a sawtooth history and a human-like recombination map. We divide samples into a group of 100 diploid samples (ref), and two groups with one diploid sample each (tsk_0 and tsk_1). Rows show the target sequence used (tsk_0 or tsk_1) and columns show the reference sequences used (ref or tsk_0), where panel tsk_0 vs tsk_0 corresponds to the within individual effective population size. For the direct MLE, we use joint trees of all 102 samples. For Colate, we use trees corresponding to the 100 diploid samples (ref) to date mutations. We also evaluate Colate in the case where true mutation ages are known.

**Supplementary Figure 3.**
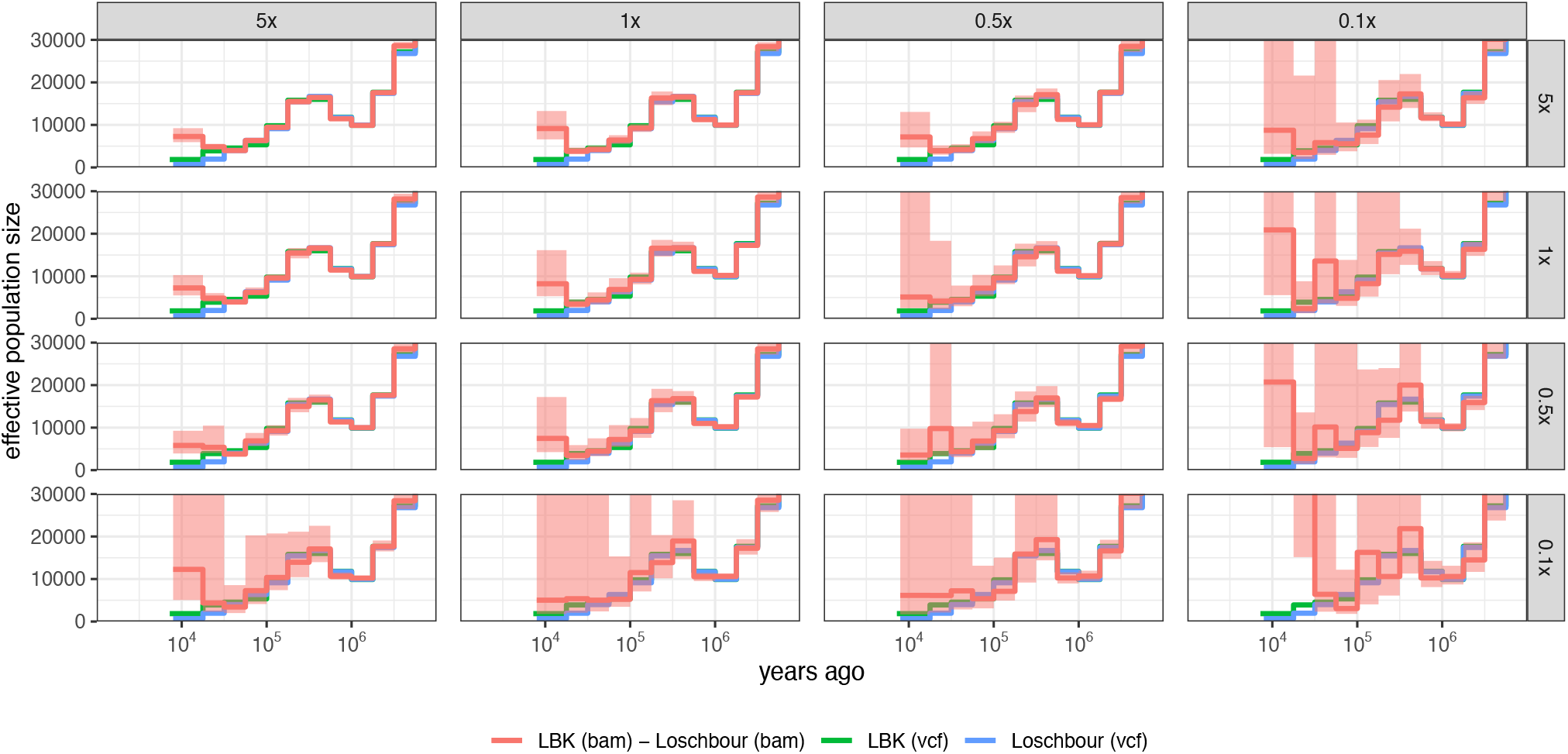
Colate-inferred effective population sizes between LBK (target sample; rows) and Loschbour (reference sample; columns), with each individual downsampled to 5x, 1x, 0.5x, and 0.1x. We additionally also show the within individual effective population sizes for each individual in green and blue, which is identical in all panels and is calculated using VCFs that were called on the original BAM files (>10x coverage).

**Supplementary Figure 4.**
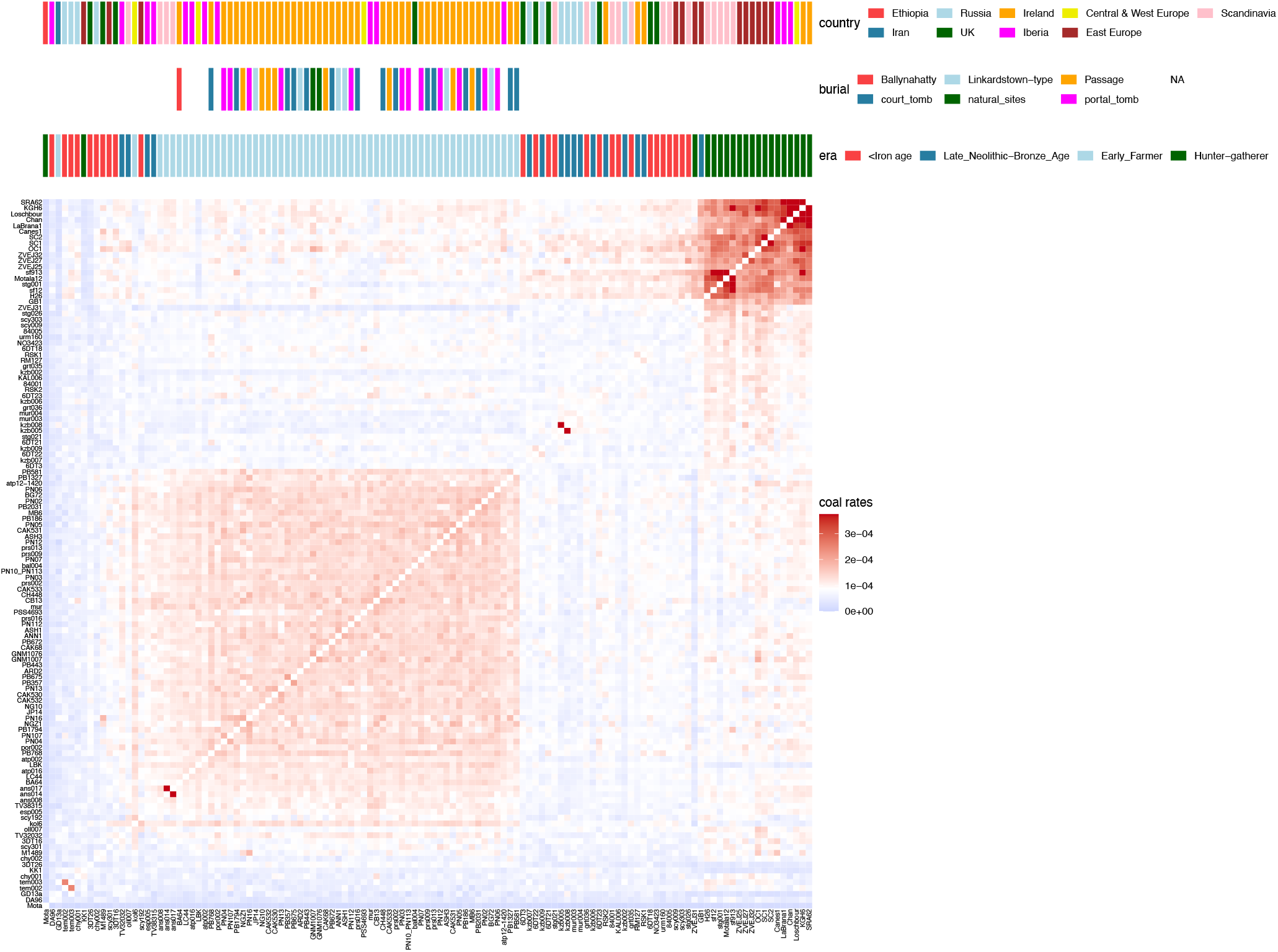
Matrix of coalescence rates calculated using Colate for epoch spanning the age of the sample to 15,000 YBP. Rows and columns of the matrix are sorted by applying UPGMA. Annotations at the top correspond to geographical region of the sample, burial type for the Irish genomes, and time period.

**Supplementary Figure 5.**
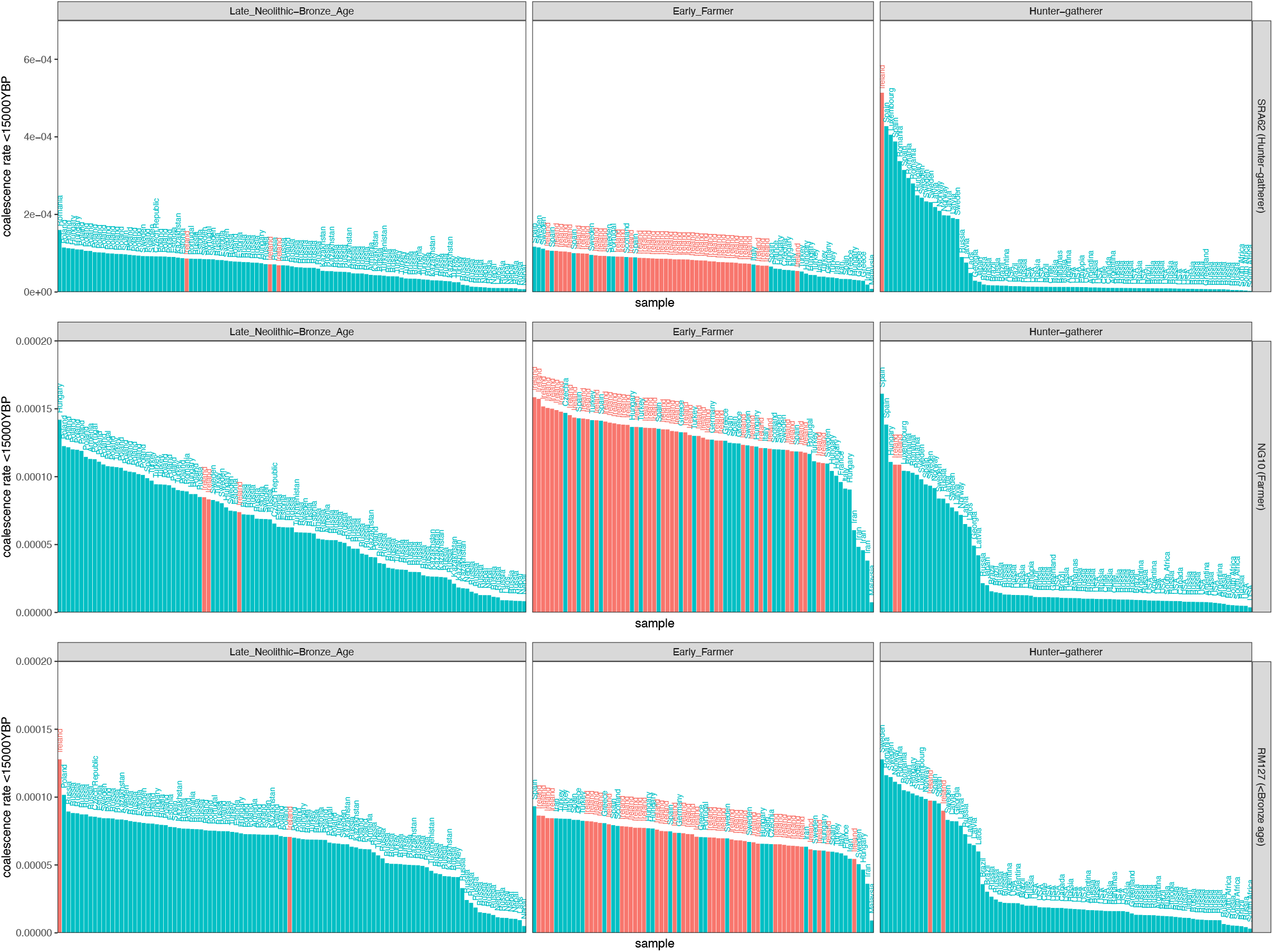
Colate-estimated coalescence rate of an Irish HG (SRA62), Irish Neolithic farmer (NG10), and an Irish Bronze-age sample (RM127) to other ancients, calculated for an epoch ranging from the date of the sample to 15,000 years BP. In each panel, samples are sorted in descending order. Colours indicates Irish samples (red) and labels annotate geographic region.

**Supplementary Figure 6.**
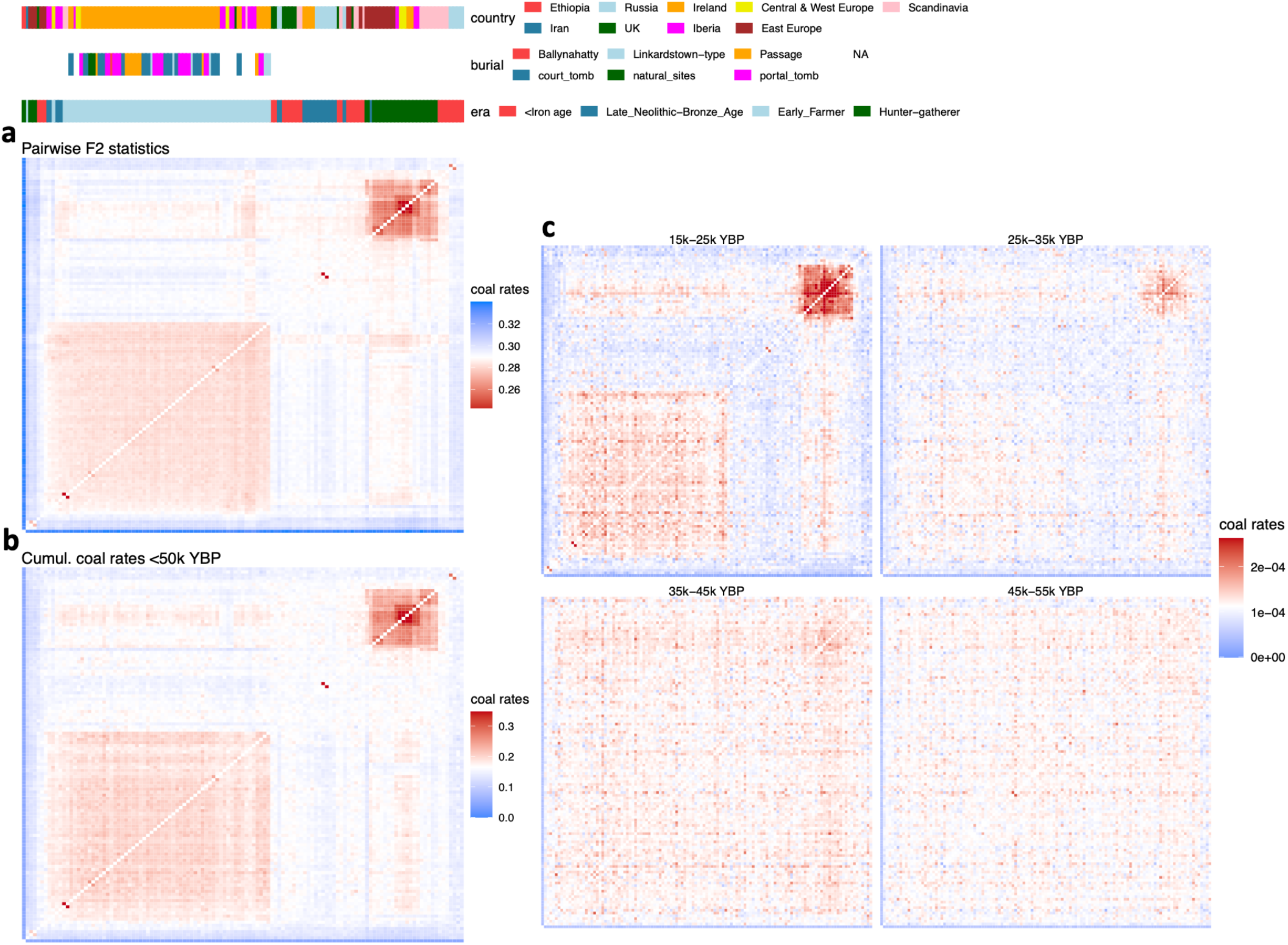
**a**, Matrix containing pairwise F2 statistics calculated using pseudohaploid calls for each individual (**Methods**). Matrix is sorted by applying UPGMA to this matrix. Annotations at the top correspond to geographical region of the sample, burial type for the Irish genomes, and time period. **b**, Matrix of Colate-inferred coalescence rates integrated over 0 - 50k YBP, ordered in the same way as the matrix in **a. c**, Matrices of pairwise coalescence rates for four epochs. All matrices are sorted in the same way as the matrix in **a**.

**Supplementary Figure 7.**
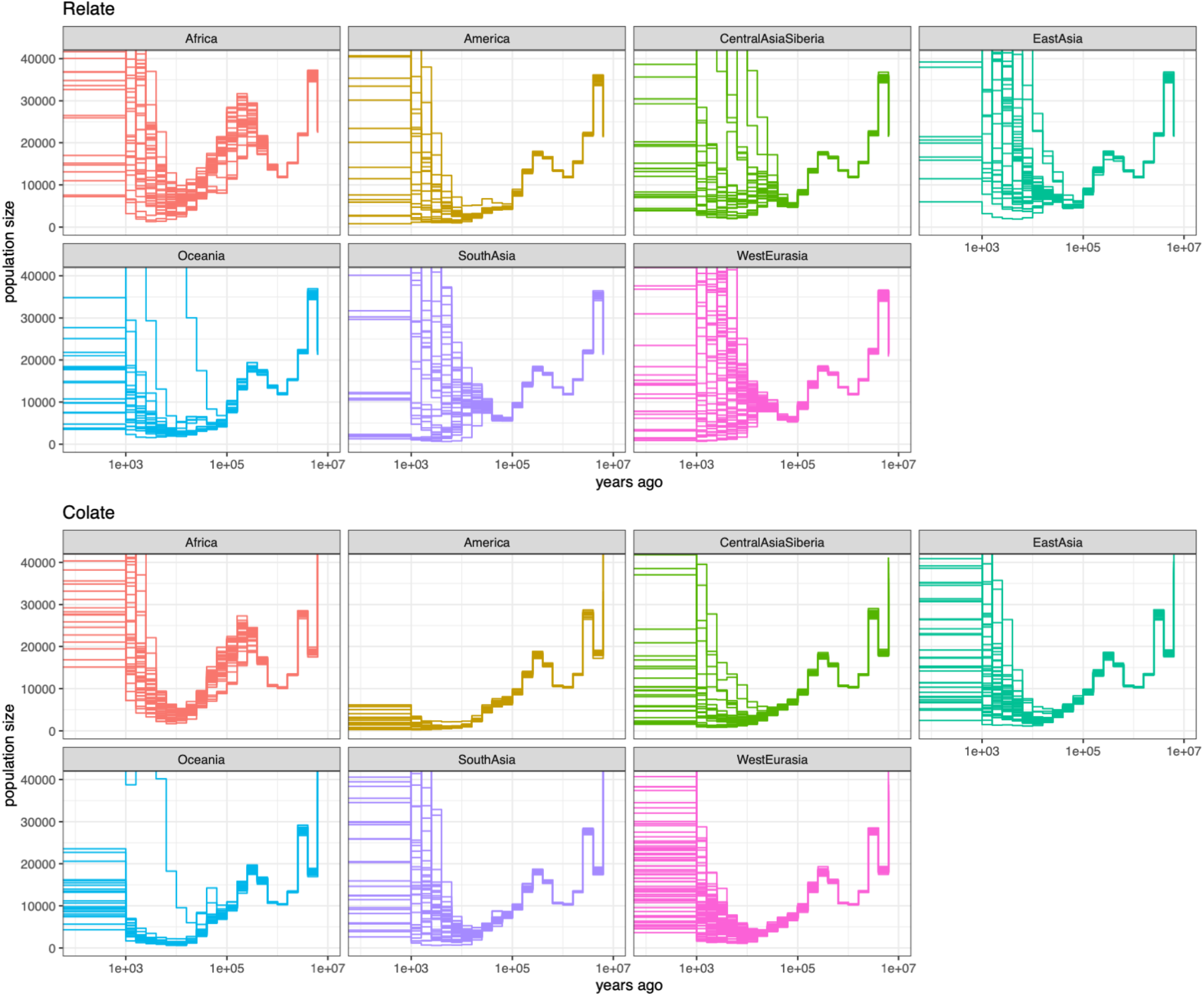
Within-individual effective population sizes for 278 samples in the Simons Genome Diversity Project inferred using *Relate* (top) and *Colate* (bottom).

**Supplementary Figure 8.**
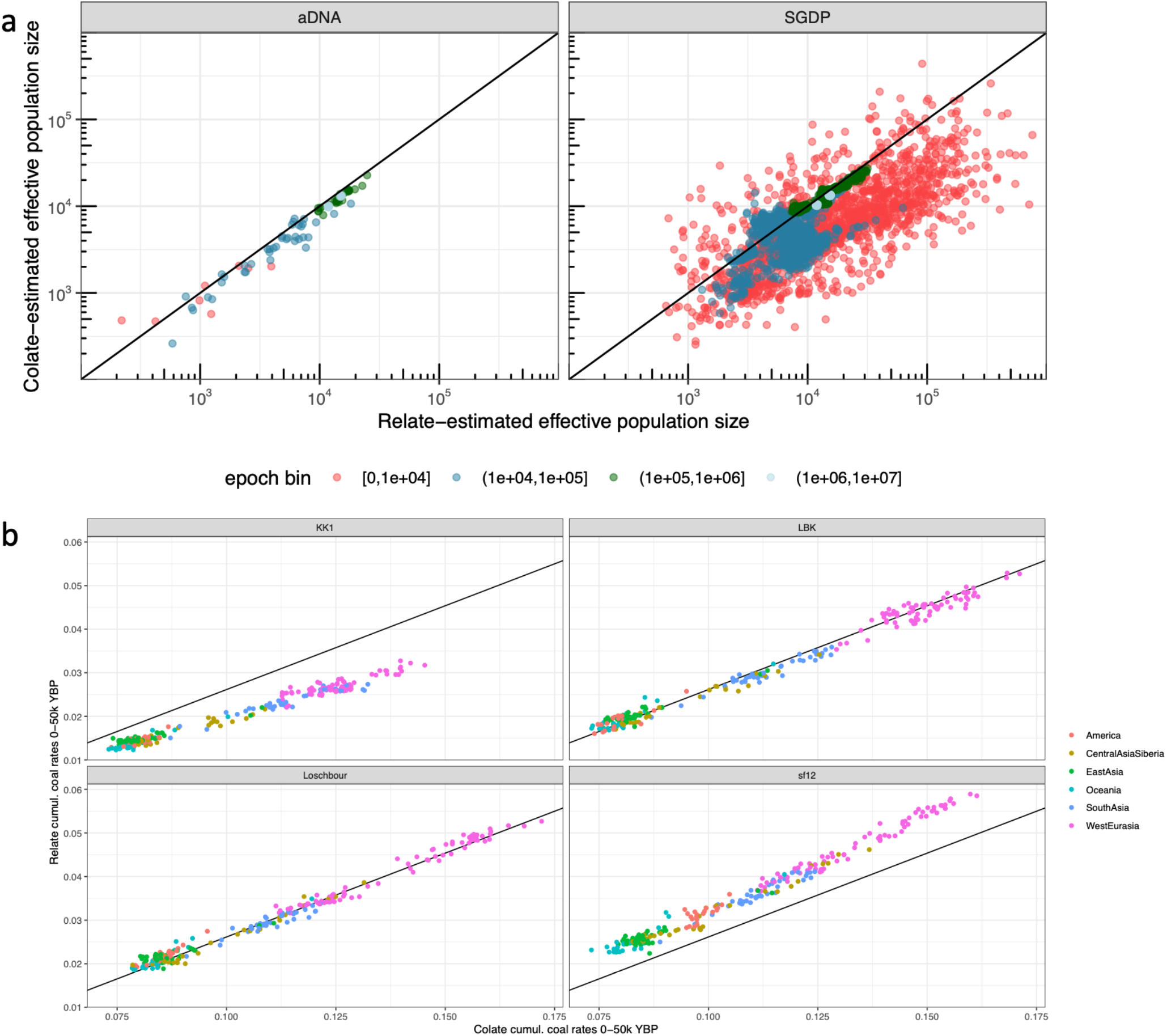
**a**, *Colate*-estimated within individual effective population sizes plotted against their *Relate*-estimated equivalents. Epochs are grouped into four bins, shown by different colours. **b**, Coalescence rates between sample shown in facet title against non-African SGDP individuals, integrated over 0 - 50k YBP, compared between *Relate* and *Colate*. We performed a linear regression on all four samples jointly, with the line shown corresponding to y = 0.38x-0.01, which was used to rescale *Colate* coalescence rates in Figure 4c.

**Supplementary Figure 9.**
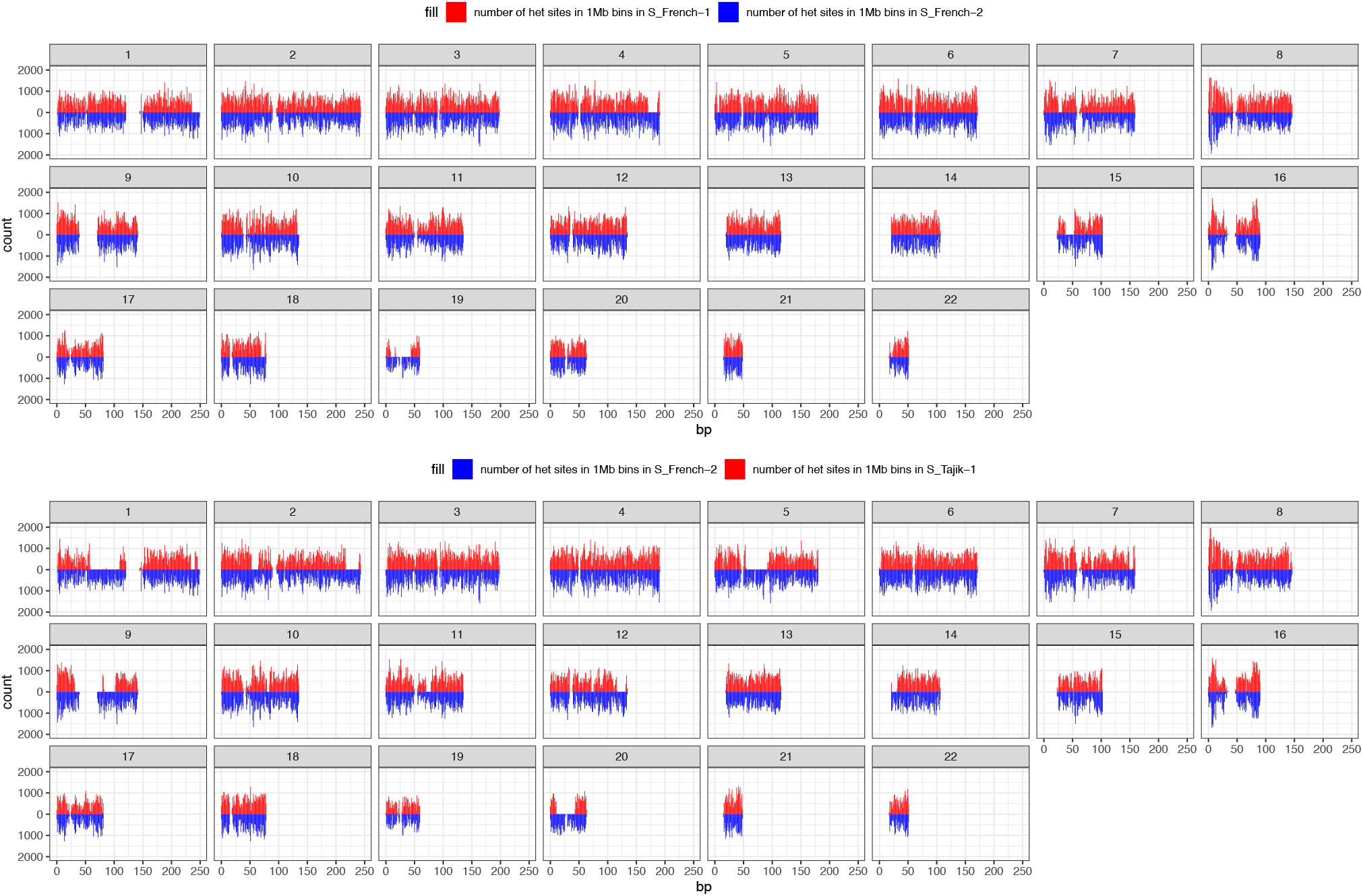
Number of heterozygous sites in 1Mb bins for the SGDP samples S_French-1 and S-Tajik-1 (red in top and bottom plot) compared to S_French-2 (blue in both plots), showing long runs of homozygosity (ROH) in S_French-1 and S_Tajik-1 compared to S_French-2. These ROH appear in different locations in S_French-1 and S_Tajik-1. While S_French-1 is a cell line, which could artificially introduce such ROH, S_Tajik-1 is a blood sample. The *Relate*-inferred effective population sizes in the most recent bin for these individuals are 10,898 for S_French-1, 161,112 for S_French-2, and 909 for S_Tajik-1.

**Supplementary Figure 10.**
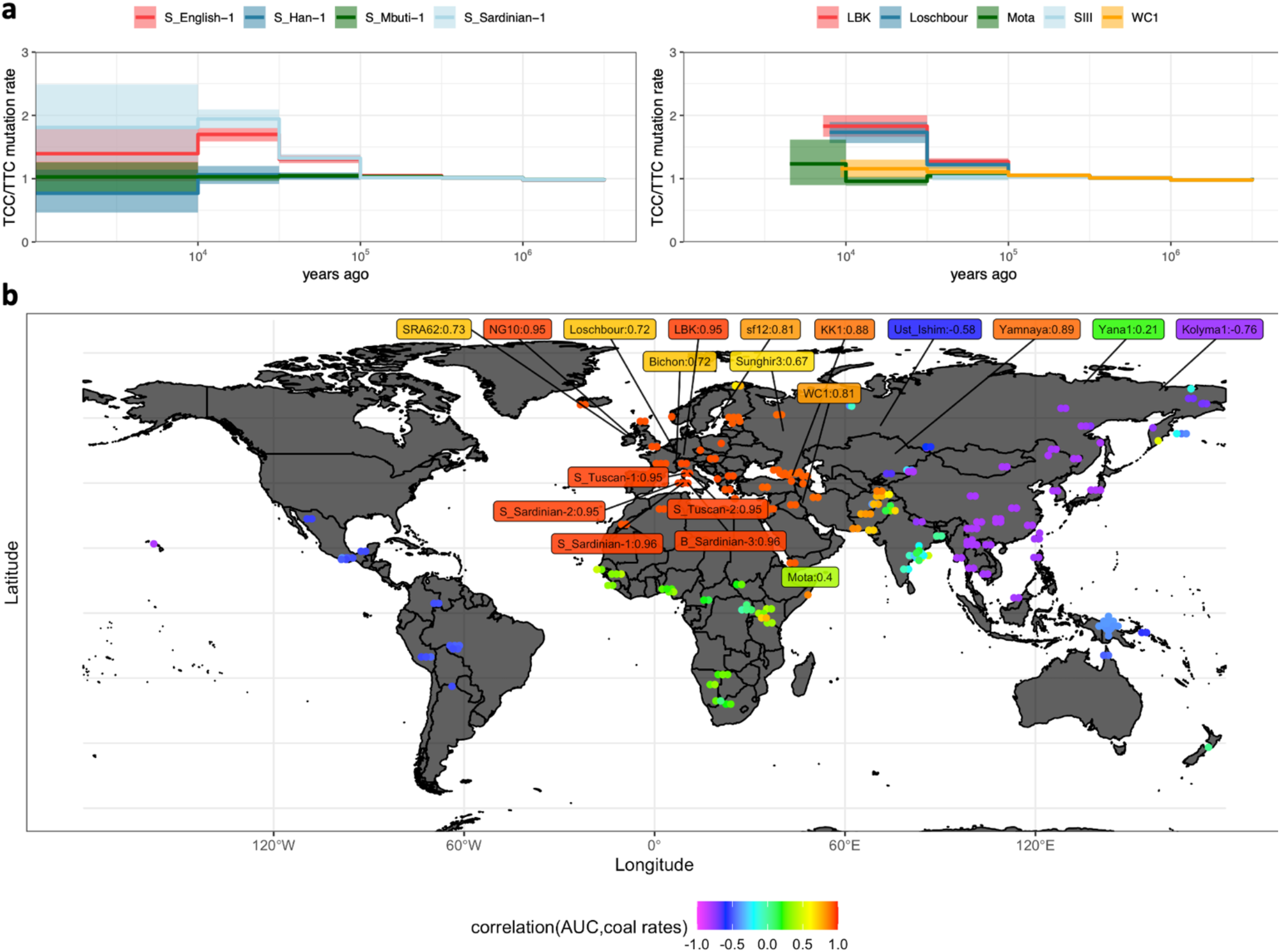
**a**, TCC/TTC mutation rate relative to the mutation rate in the time interval 100k-1M YBP for four modern individuals and five ancient individuals. **b**, Correlation calculated between the “area under the curve” (AUC) of the TCC/TTC mutation rate (**Methods**) and Colate-inferred coalescence rates to all non-African SGDP individuals and non-Africans ancients. Correlations for SGDP individuals are shown by circles and correlations for ancient individuals are labelled.

**Supplementary Figure 11.**
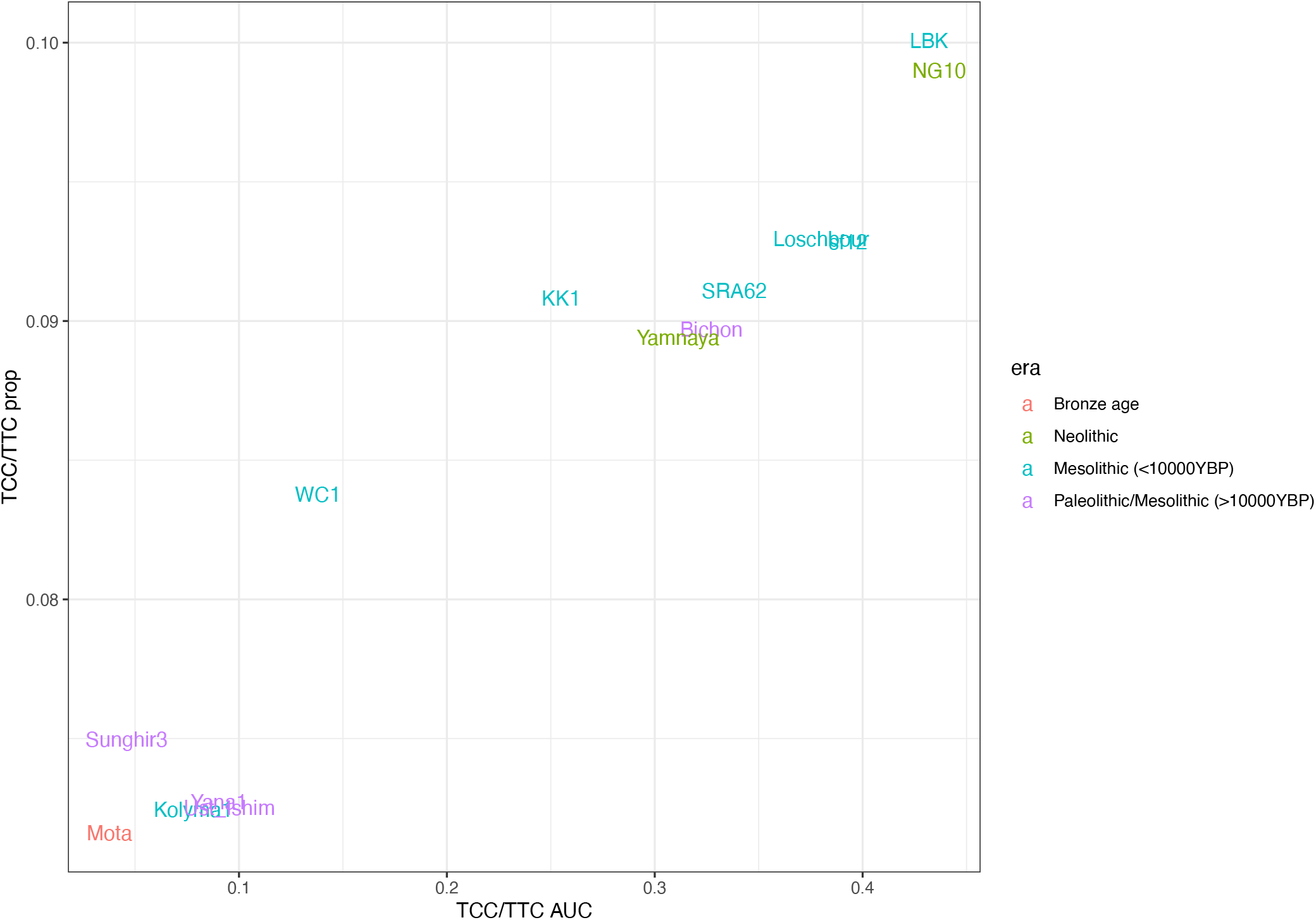
Comparison of two different ways of quantifying the TCC/TTC mutation rate signature plotted against each other (**Methods**). X-axis shows area under the curve (AUC) calculated from mutation rates directly obtained using *Relate* genealogies, whereas y-axis shows the number of TCC/TTC mutations relative to other transitions (excl. CpGs), for mutations dated to be younger than 100k YBP.

**Supplementary Figure 12.**
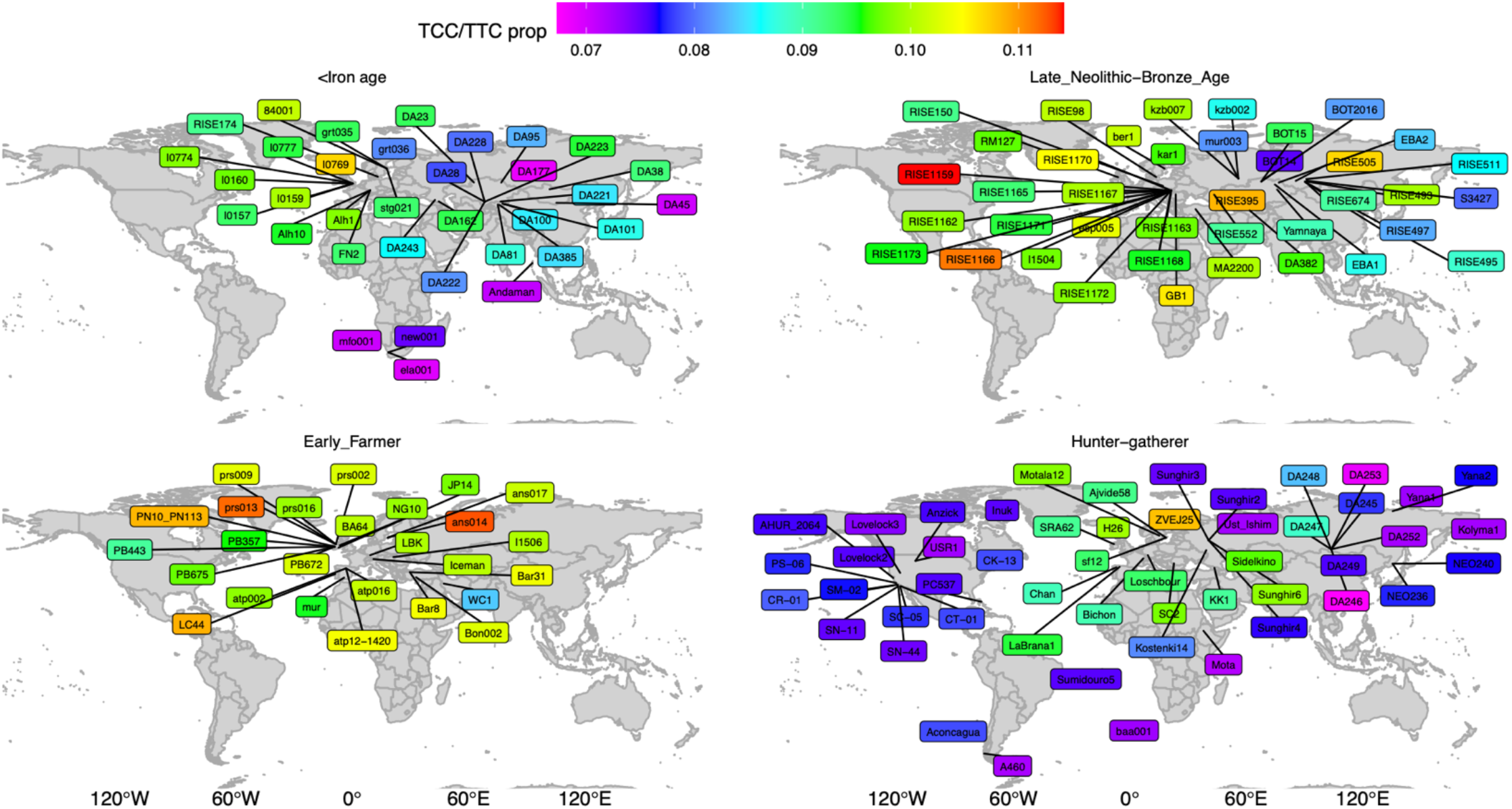
The proportion of TCC/TTC mutations relative to C/T transitions (excluding those in CpG contexts) (**Methods**), for ancient samples of >2x mean coverage. Each map shows a different cultural context/time period and colours indicate signal strength.

**Supplementary Figure 13.**
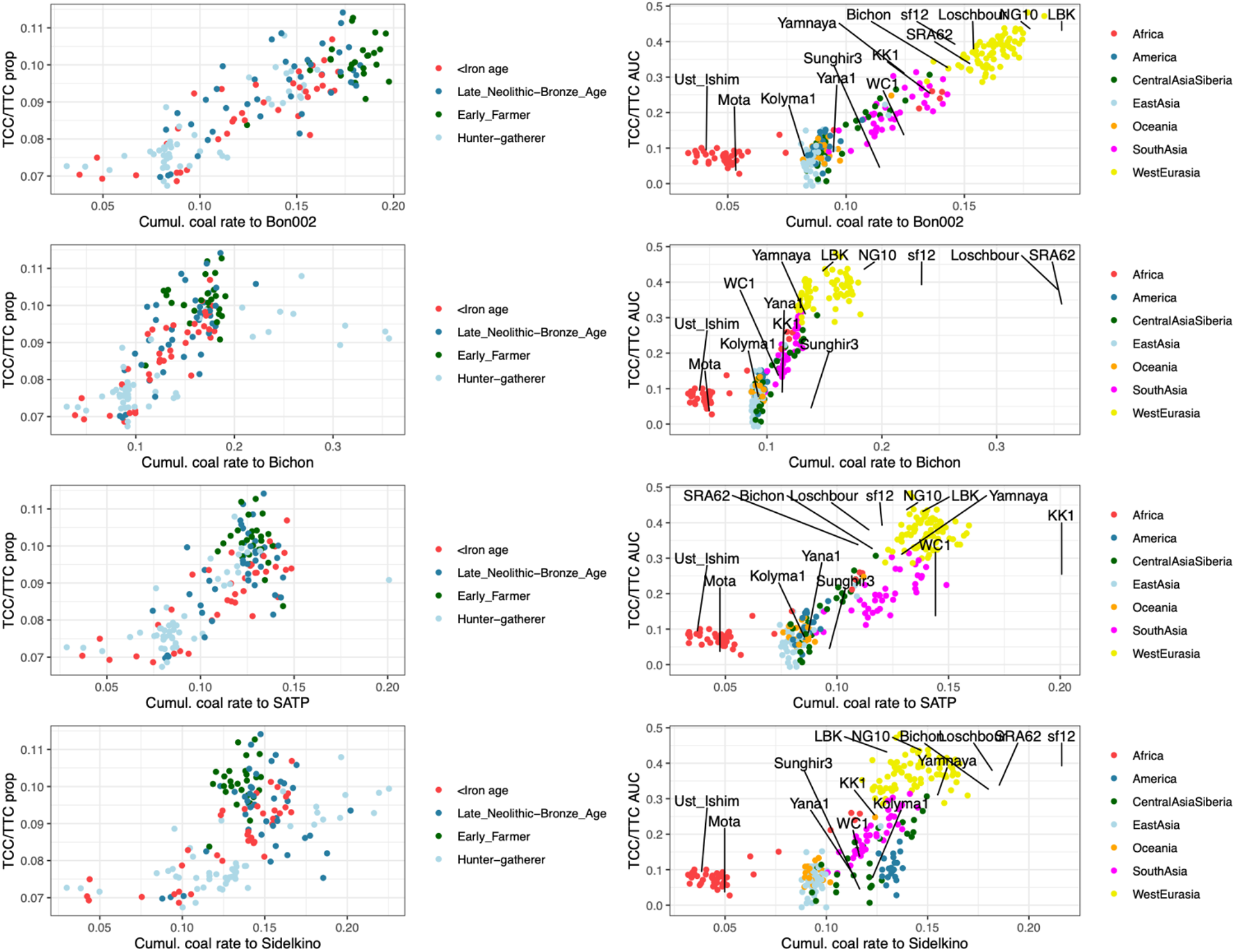
Strength of the TCC/TTC mutation rate signal, quantified using the proportion of TCC/TTC mutations relative to transitions (left column) or area under the mutation rate curve (right column) (**Methods**) plotted against cumulative coalescence rates with Bon002, a 10k-year-old Anatolian individual, Bichon, a 13k-year-old Western HG, SATP, a 13k-year-old Caucasus HG, and Sidelkino, a 11k-year-old Eastern HG. The cumulative coalescence rates are calculated as the integral of the coalescence rate from sample age to 50k YBP.

**Supplementary Figure 14.**
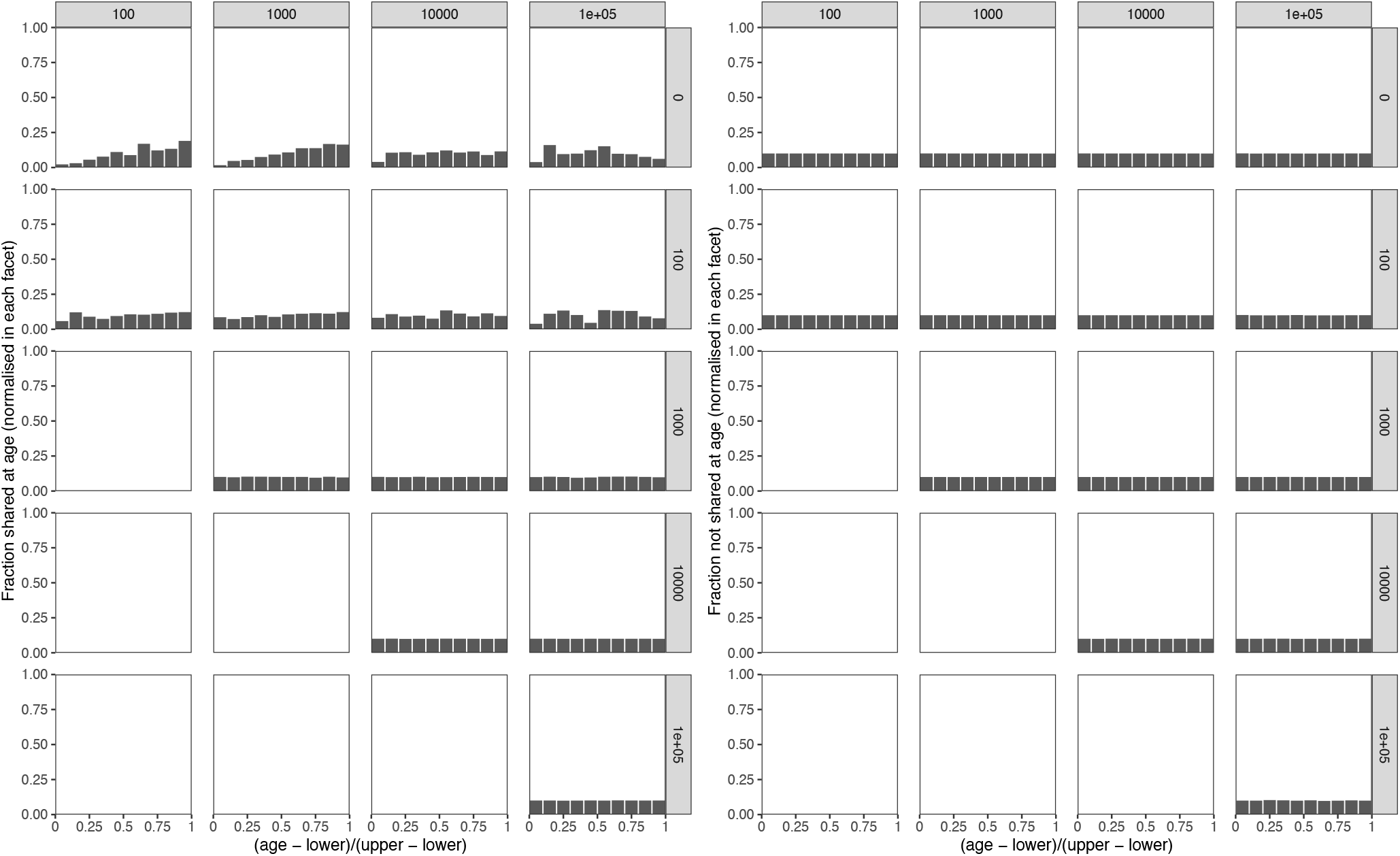
For mutations segregating in the 100 diploid samples (ref) in the zigzag simulation of Supplementary Figure 2, we plot a histogram of the true age of the mutation relative to lower and upper ages of the coalescence events of the branch on which this mutation occurred, using the genealogy of these 100 diploid samples only and stratified by whether or not it is shared with sample tsk_0 (left and right panel). We additionally stratify by age bins of lower (rows) and upper (columns) coalescence ages. This shows that mutations that are singletons in the group of 100 diploid samples and are shared with tsk_0 have a non-uniform age, whereas all other categories are close or nearly identical to uniform distributions.

